# Relaxin-2 drives regenerative healing and suppresses scar formation

**DOI:** 10.64898/2025.12.08.692836

**Authors:** Amanda K. Williamson, Katherine M. Hohl, Jack R. Kirsch, Rosalynn M. Nazarian, Thomas P. Schaer, Daniel S. Roh, Mark W. Grinstaff

**Affiliations:** Department of Chemistry, Boston University, Boston, MA, 02215, USA; Department of Biomedical Engineering, Boston University, Boston, MA, 02215, USA; Pathology Service, Dermatopathology Unit, Massachusetts General Hospital, Harvard Medical School, Boston, MA 02114, USA; Penn Vet Institute for Medical Translation, University of Pennsylvania School of Veterinary Medicine New Bolton Center, Kennett Square, PA, 19348, USA; Division of Plastic and Reconstructive Surgery, Department of Surgery, Boston University School of Medicine, Boston, MA, 02118, USA

**Keywords:** Scars, Fibrosis, TGF-β1-Smad pathway, Relaxin-2

## Abstract

Fibrotic scarring is a pervasive and unresolved challenge in medicine, leading to permanent disfigurement, impaired mobility, and severe disruption of basic skin functions including elasticity, barrier protection, and thermoregulation. Despite its far-reaching personal, clinical, and economic impact, affecting hundreds of millions worldwide after surgery, trauma, and burns, no effective treatments exist to halt or reverse pathological scar formation. Scarring results from uncontrolled TGF-b1 signaling, which drives excessive deposition of extracellular matrix (ECM) proteins such as collagen-I/III and accumulation of alpha-smooth muscle actin (alpha-SMA), producing rigid, dysfunctional tissue. Here, we present a mechanistically guided approach targeting this unmet clinical need, leveraging the natural antifibrotic peptide hormone relaxin-2 (RLX-2) to actively remodel dermal architecture. RLX-2 signals via its G-protein coupled receptor RXFP1, upregulating matrix metalloproteinases (MMPs) and inhibiting aberrant ECM production. In TGF-b1-activated dermal fibroblasts across 2D and 3D in vitro models, ex vivo healthy and scarred human skin samples - cultured under physiological and pathological tension - and in an in vivo murine burn wound model, RLX-2 robustly suppresses fibrosis, restores regenerative tissue features, and rescues dermal architecture. Importantly, RLX-2 achieves this result without compromising the normal wound healing process, highlighting its potential as a transformative therapy for both prevention and reversal of pathological scarring.

## Introduction

Scarring arises due to aberrant wound healing in response to traumatic injury to the skin. Hypertrophic scars are common, developing in 91% of patients with burn injuries and in 40-70% of post-surgical patients, as well as following acne, vaccination, and infection^1^. Mechanistically, a prolonged inflammatory state characterized by a cascade of cytokines and growth factors, including transforming growth factor-β1 (TGF-β1), triggers the excessive deposition of extracellular matrix (ECM), the defining characteristic of scar tissue. When unchecked, TGF-β1 signaling drives uncontrolled differentiation of fibroblasts into myofibroblasts, eliciting overproduction of collagen-I/III, α-smooth muscle actin (αSMA), and fibronectin and altering ECM homeostasis via downregulation of collagen-degrading matrix metalloproteinases (MMPs) and upregulation of tissue inhibitors of metalloproteinases (TIMPs)^2,3^. At the tissue scale, these changes afford large, stiff scars that are frequently painful, pruritic, mechanically constrictive, disfiguring, and psychologically detrimental^4–8^. Morphologically, hypertrophic scars possess dense, highly aligned collagen fibers that orient parallel to the epidermis, remain within the bounds of the original wound, and lack key dermal appendages such as hair follicles and sweat glands. This altered structure yields diminished mechanical strength and compromised function, affecting barrier integrity, elasticity, and thermoregulation^7,9–11^.

Despite significant scientific effort, effective strategies for scar prevention and reversal remain elusive. Current treatments include surgery, corticosteroid injections, radiotherapy, laser therapy, cryotherapy, compression, and cytotoxic drugs, or a combination thereof; however, these therapeutic approaches are still only partially successful and are hindered by high recurrence rates and adverse effects^5,8,10,12^. Consequently, we hypothesized that a tissue remodeling therapeutic strategy will offer significant benefits over existing treatments that merely manage symptoms and/or rely on tissue ablation. Apart from scarless wound healing observed during fetal development, very few tissues in the body undergo regeneration after a significant insult; however, native biologic mechanisms that facilitate ECM remodeling provide insight into potential signaling pathways to leverage. For example, during pregnancy, the endogenous peptide hormone relaxin-2 (RLX-2) drives remodeling of uterine, cervical and birth canal ECM via inhibition of TGF-β1 signaling to facilitate embryo implantation, placental development, and delivery as well as stimulates a decrease in systemic vascular resistance, particularly in the uterine circulation, to support the needs of the growing fetus^13^. After birth, the altered tissues return to a normal, healthy structure and function. As a result of these beneficial effects, RLX-2 is of interest as an antifibrotic and vasodilatory agent for pathologies involving the heart, kidney, liver, lung, and joint tissues^14–20^. With its natural ability to target the underlying pathology of fibrosis via inhibition of the pro-fibrotic TGF-β1 pathway, we propose RLX-2 a promising therapeutic for regenerative, scarless wound healing.

Herein, we report that RLX-2 effectively inhibits scar formation and promotes regenerative wound healing across models of increasing complexity, including *in vitro*, *ex vivo,* and *in vivo* systems. Tracking transcriptional and translational expression changes in both 2D and 3D *in vitro* models of dermal fibrosis reveals that RLX-2 modulates key components of the TGF-β1 signaling pathway, underlying its mechanism of action. In human skin tissue cultured *ex vivo*—both with and without tension as a multicellular, three-dimensional model of scar induction— RLX-2 prevents fibrosis, restores healthy dermal and epidermal tissue, and rescues normal tissue architecture. In an *in vivo* murine chemical burn wound model, RLX-2 accelerates wound healing and significantly reduces scar area. Finally, in skin samples from patients with hypertrophic scars, RLX-2 demonstrates promising regenerative capacity, suggesting its potential as a therapeutic strategy for both scar prevention and treatment of clinically significant scars.

## Results

### RLX-2 suppresses TGF-β1-induced pro-fibrotic phenotype in human dermal fibroblasts

RLX-2 signals via its cognate G-protein coupled receptor, RXFP1^21–23^. Activation of RXFP1 signaling by RLX-2 in primary human dermal fibroblasts leads to dose dependent increases in intracellular cAMP concentration (**Figure 1A, i**), indicating receptor engagement. Co-treatment with forskolin, an adenylyl cyclase activator, produces no further change in cAMP with RLX-2, confirming selective activation of the stimulatory Gαs subunit and excluding inhibitory Gαi signaling (**Figure 1A, ii**)^24^.

**Figure 1.**
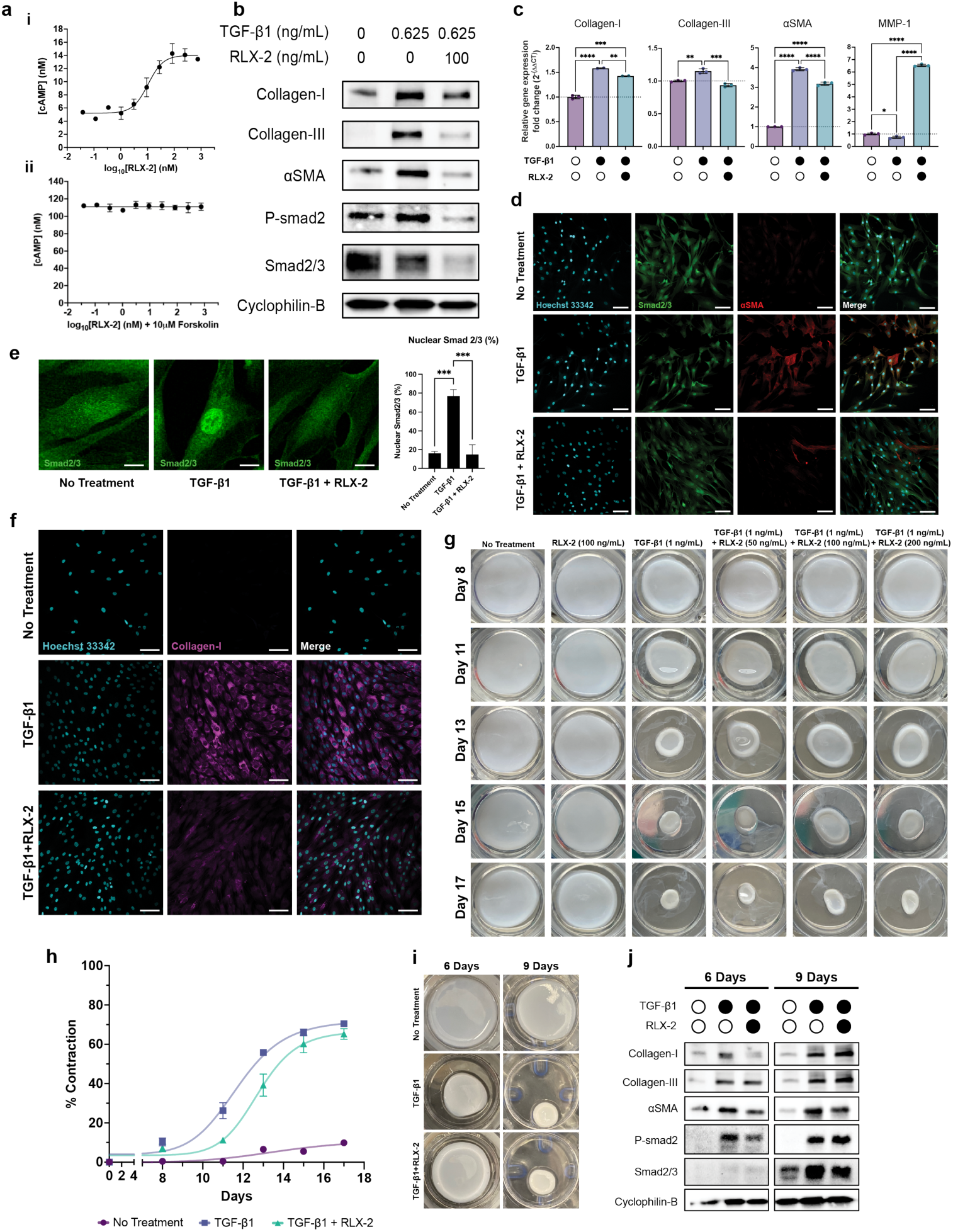
*In vitro* characterization of RLX-2 antifibrotic effects in normal human dermal fibroblasts. (**A**) cAMP concentration measured in human dermal fibroblasts (EC50 = 10.5 nM = 62.62 ng/mL) treated with increasing concentrations of RLX-2 only (i) and with 10 μM forskolin (ii). (**B**) Protein expression in normal dermal fibroblasts treated with TGF-β1 (0.625 ng/mL) and RLX-2 (100 ng/mL) for 24 hours. (**C**) RT-qPCR for gene expression of collagen-I, collagen-III, αSMA and MMP-1 in normal dermal fibroblasts treated with TGF-β1 (0.625 ng/mL) with and without RLX-2 (100 ng/mL) for 24 hours. Relative expression level is compared to GAPDH as the reference gene and the untreated control. Error bars show standard deviation of technical replicates. Statistical significance was determined by an ordinary one-way ANOVA using a Tukey test to control for multiple comparisons. *=p<0.05, **=p<0.005, ***=p<0.0005, ****=p<0.0001. (**D**) Immunostaining of smad2/3 and αSMA in normal dermal fibroblasts treated with TGF-β1 (0.2 ng/mL) and RLX-2 (100 ng/mL) for 24 hours. Cyan = nucleus, Green = smad2/3, Red = αSMA. 40X magnification, scale bar = 100μM (**E**) Images of smad2/3 immunostaining and quantification of smad2/3 nuclear partitioning. Scale bar = 15μM (**F**) Immunostaining of collagen-I in normal dermal fibroblasts treated with TGF-β1 (5 ng/mL) and RLX-2 (100 ng/mL) for 3 days. Cyan = nucleus, Magenta = collagen-I. 20X magnification, scale bar = 100μM (**G)** Contraction of 3D collagen gels embedded with dermal fibroblasts and treated with TGF-β1 (1 ng/mL) with and without increasing doses of RLX-2 (50 – 200 ng/mL) over 17 days. (**H**) Gel contraction was calculated using the change in diameters compared to the original dimensions of the transwell. Media and treatments at indicated doses were changed every 2 – 3 days (n=2). (**I**) Visualization of collagen gel contraction by normal dermal fibroblasts treated with TGF-β1 (1 ng/mL) with and without RLX-2 (200 ng/mL). (**J**) Corresponding expression of fibrotic proteins and P-smad2 at 6 and 9 days. All data in this figure is representative of trends observed in at least biological duplicate.

In the canonical TGF-β1 pathway, binding of TGF-β1 to its receptor causes phosphorylation of Smad2 and Smad3, which translocate to the nucleus with Smad4 to drive transcription of ECM proteins such as collagen-I and collagen-III and intracellular proteins such as αSMA^25^. TGF-β1 stimulation induces robust myofibroblast differentiation, as evidenced by nuclear translocation of Smad2/3 and upregulation of ECM components such as collagen-I, collagen-III, and αSMA (**Figure 1B-F**). Co-treatment with RLX-2 reverses nuclear Smad2/3 accumulation over 24 hours (**Figure 1D-E**). This reduction of nuclear Smad2/3 also corresponds to a decrease in αSMA, supporting that the antifibrotic effects of RLX-2 occur through TGF-β1-Smad pathway inhibition and abrogate myofibroblast differentiation in dermal fibroblasts (**Figure 1D**). Furthermore, RLX-2 reduces the prevalence of P-Smad2, suggesting that RLX-2 blocks TGF-β1-mediated Smad2/3 phosphorylation as the mechanism for reduction of Smad2/3 nuclear accumulation and subsequent suppression of downstream TGF-β1 pathway effects (**Figure 1B**).

RLX-2 co-treatment attenuates TGF-β1-stimulated increases in collagen-I, collagen-III, and αSMA at the transcriptional and translational levels, indicating robust suppression of the pro-fibrotic phenotype (**Figure 1B-D, F**). RLX-2 also restores MMP-1 gene expression following TGF-β1-induced downregulation, supporting a dual impact of RLX-2 on limiting matrix accumulation and promoting ECM degradation (**Figure 1C**).

### RLX-2 delays early collagen matrix contraction in 3D in vitro fibroblast model

It is known that fibroblasts grown on stiff plastic cell culture plates experience unnaturally high mechanical forces that promote a fibrotic phenotype, and that cells more generally modulate their phenotype based on substrate stiffness^26–28^. To better recapitulate the soft, three-dimensional environment that fibroblasts inhabit *in vivo*, we embedded dermal fibroblasts in type I collagen gels, treated with TGF-β1 and RLX-2, and observed the rate of gel compaction. Because increased tensile forces and tissue contraction are major contributors to the formation of hypertrophic scars^29–31^, this 3D collagen gel model offers a strategy to observe changes in bulk contraction in response to TGF-β1 and RLX-2^32,33^. Fibroblast-driven contraction of type I collagen gels, normally accelerated by TGF-β1, delays in the presence of RLX-2 co-treatment across a 17-day interval (**Figure 1G-H**). RLX-2 does not fully abrogate contraction but significantly reduces its rate during the initial 10 days. Expression of matrix proteins and P-Smad2 remain elevated in TGF-β1 conditions, while RLX-2 co-treatment most strongly curtails the pro-fibrotic phenotype at early timepoints, coinciding with maximal inhibition of gel contraction (**Figure 1I-J**). By Day 9, both groups exhibit comparable contraction and ECM protein expression. These findings indicate that RLX-2 selectively inhibits excessive early dermal contraction without impairing basal contractility necessary for normal wound closure.

### RLX-2 promotes regenerative phenotype in ex vivo human skin models

Building on insights gained from 2D and 3D systems, we interrogated how RLX-2’s antifibrotic effects operate within native *ex vivo* tissue architecture. Thus, we examined the effects of RLX-2 on human skin samples cultured *ex vivo* to evaluate the ability of RLX-2 to prevent a TGF-β1-induced scar phenotype (**Figure 2A**). TGF-β1 treatment (1 ng/mL) of *ex vivo* human skin biopsies (Patient 1) induces dose-dependent upregulation of collagen-I, collagen-III, αSMA, CTGF, and TIMP-1, indicative of a pro-fibrotic state. RLX-2 at 100 ng/mL effectively restores these transcripts to near-basal levels, while the 200 ng/mL dose preferentially increases MMP-1/9 expression, shifting the response toward enhanced ECM remodeling (**Figure 2B, S1B**). At the protein level, TGF-β1 (2 ng/mL) induced upregulation of collagen-I, collagen-III, and fibronectin returns to the untreated control level with RLX-2 treatment (100 ng/mL). A higher dose of RLX-2 (200 ng/mL) only partially restores collagen-I and fibronectin levels, with minimal effect on collagen-III (**Figure 2C**). RLX-2 substantially increases αSMA protein at 100 ng/mL, independent of transcript levels (**Figure 2B-C**). Of note is that treatment with 1 ng/mL TGF-β1 does not show significant increases in pro-fibrotic ECM proteins at the translational level, despite increases observed at the transcriptional level (**Figure S1A-S1B**).

**Figure 2.**
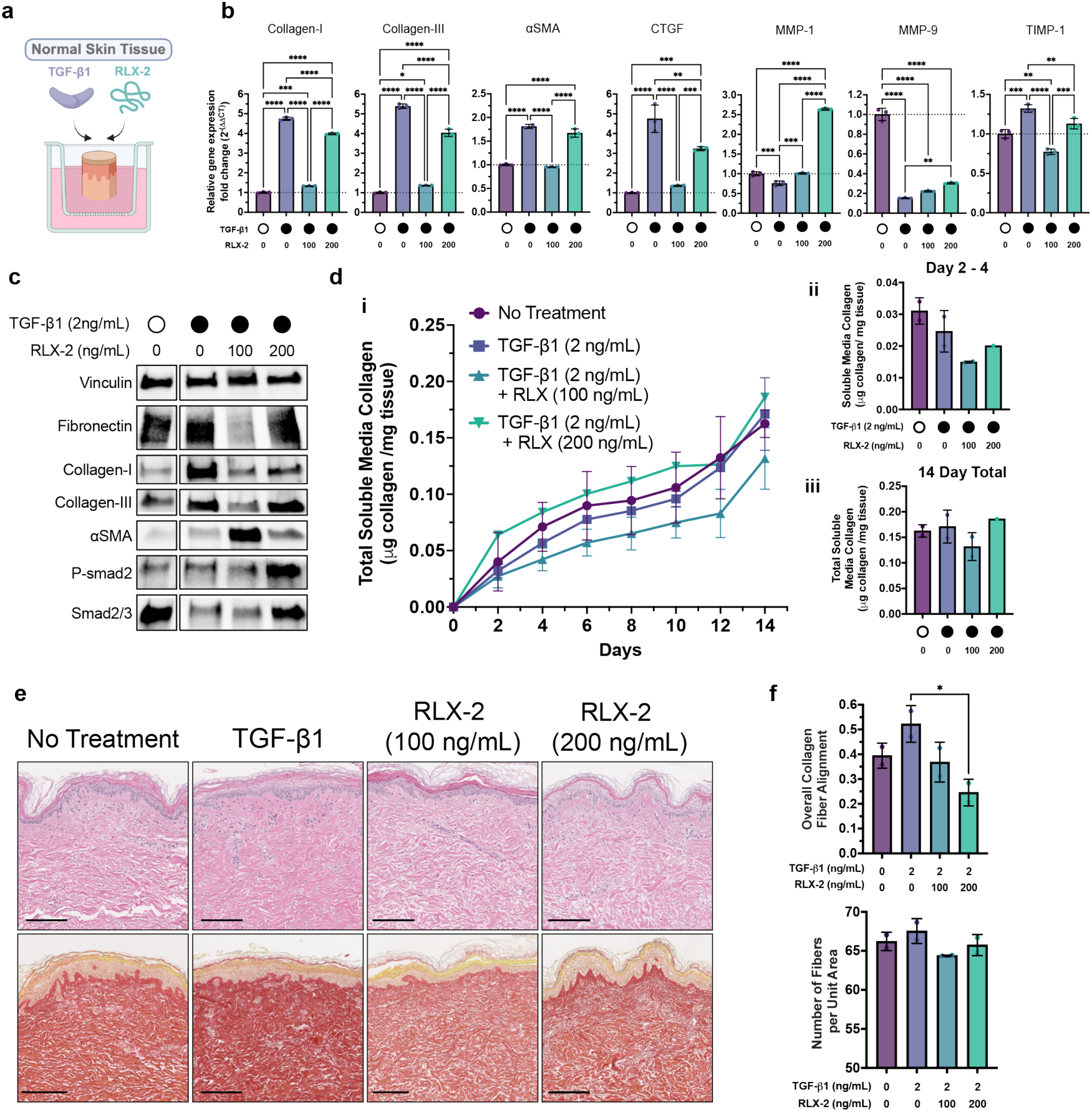
RLX-2 inhibits the fibrotic scar phenotype in ex vivo normal human skin tissues. (**A**) 8 mm biopsy punches were made in normal human skin tissue samples and cultured at an air-liquid interface with TGF-β1 and RLX-2 treatments added to the media for 14 days. All data shown is using tissue from Patient 1 cultured *ex vivo* with TGF-β (2 ng/mL) with and without RLX-2 (100 or 200 ng/mL) except in (F). (**B**) RT-qPCR of collagen-I, collagen-III, αSMA, CTGF, MMP-1, MMP-9, TIMP-1. (**C**) Protein expression of ECM proteins, αSMA, Smad2/3, and P-smad2. (**D**) (i) Cumulative soluble collagen released in the media every two days over 14 days. (ii) Soluble collagen released in media between treatment days 2 and 4. (iii) Total soluble collagen released after 14 days. Collagen content was normalized to the total mass of the biopsy punch. Bars show range of biological duplicates (n=2 biopsy punches from Patient 1). (**E**) Representative images of H&E and Sirius red staining of normal human skin tissues after 14 days of *ex vivo* culture. Top = H&E, Bottom = Sirius red. Scale bar = 200 μm. (**F**) CurveAlign analysis on overall fiber alignment from Sirius red stained images of skin tissues. Points represent average of analysis performed on 8 ROIs per tissue section. Bars show range of averages from 2 different patient samples/experiments (n=8 ROIs, N=2 patients). Statistical significance was determined by an ordinary one-way ANOVA using a Tukey test to control for multiple comparisons. *=p<0.05, **=p<0.005, ****=p<0.0001.

Next, we treated the *ex vivo* skin tissue (Patient 2) with increasing TGF-β1 doses (1, 2, and 10 ng/mL) to further examine the development and reversal of this fibrotic phenotype *ex vivo*. Similarly to the Patient 1 sample, the Patient 2 sample exhibits a minimal fibrotic response to treatment with 1 ng/mL TGF-β1, particularly at the protein level (**Figure S2A**). Interestingly, at 2 ng/mL TGF-β1, while fibronectin protein levels increase significantly, none of the other markers of TGF-β1 induced fibrosis increase in comparison to the untreated control. However, at the highest dose of TGF-β1 (10 ng/mL), collagen-I, collagen-III, and fibronectin all significantly increase, both at the transcriptional and translational levels (**Figure S2A, S3C**), similarly to the increases observed with the Patient 1 sample when treated with 2 ng/mL TGF-β1. Subsequent RLX-2 treatment reveals several observations. At the highest 10 ng/mL TGF-β1 dose, RLX-2 affords the greatest reduction in ECM production with the corresponding higher 200 ng/mL dose (**Figure S2A, S3C**). Interestingly, for the 2 ng/mL TGF-β1 treatment sample, the 100 ng/mL RLX-2 dose increases levels of collagen-III at both the transcriptional and translational levels (**Figure S2A, S3B**), and at 10 ng/mL TGF-β1, neither dose of RLX-2 alters collagen-III levels significantly from the levels of the skin treated with TGF-β1 only. As increased collagen-III is a hallmark of scarless fetal skin ^34^, these findings may point to RLX-2 driving the ECM toward a regenerative, scarless phenotype.

RLX-2 treatment reduces TGF-β1-stimulated increases in soluble collagen present in culture media at early timepoints, but total collagen at endpoint remains unaffected, indicating time-restricted inhibition of excess matrix deposition (**Figure 2D, S1C**).

Collagen fibers in normal skin exhibit a haphazard basket-weave-like structure with a relatively low fiber alignment. Histological analysis of *ex vivo* treated skin from both patients reveals that TGF-β1 increases collagen alignment and reduces rete ridge presence, typical histologic findings in hypertrophic scars^35^. RLX-2 consistently reduces fiber alignment and restores rete ridges, enhancing tissue architecture resemblance to normal skin in samples from both patients (**Figure 2E-F, S1D-E, S2B-C**).

### RLX-2 rescues skin architecture from mechanical stress-induced fibrosis in ex vivo human skin

We next tested if RLX-2 treatment inhibits a fibrotic phenotype in *ex vivo* normal human abdominal tissue placed in a tension-generating device to replicate the native mechanical environment of skin (TenSkin^TM^ model; **Figure 3A**). Mechanical tension is a critical driver of fibrotic scar formation, with hypertrophic scars most commonly arising in areas of the body exposed to sustained tensile forces, such as the chest, shoulders, feet, hands, and joints^10,31,36,37^. Mechanistically, these forces induce integrin-mediated signaling cascades, including liberation of latent TGF-β1 from the ECM, which further exacerbate the fibrotic phenotype^38–40^.

**Figure 3.**
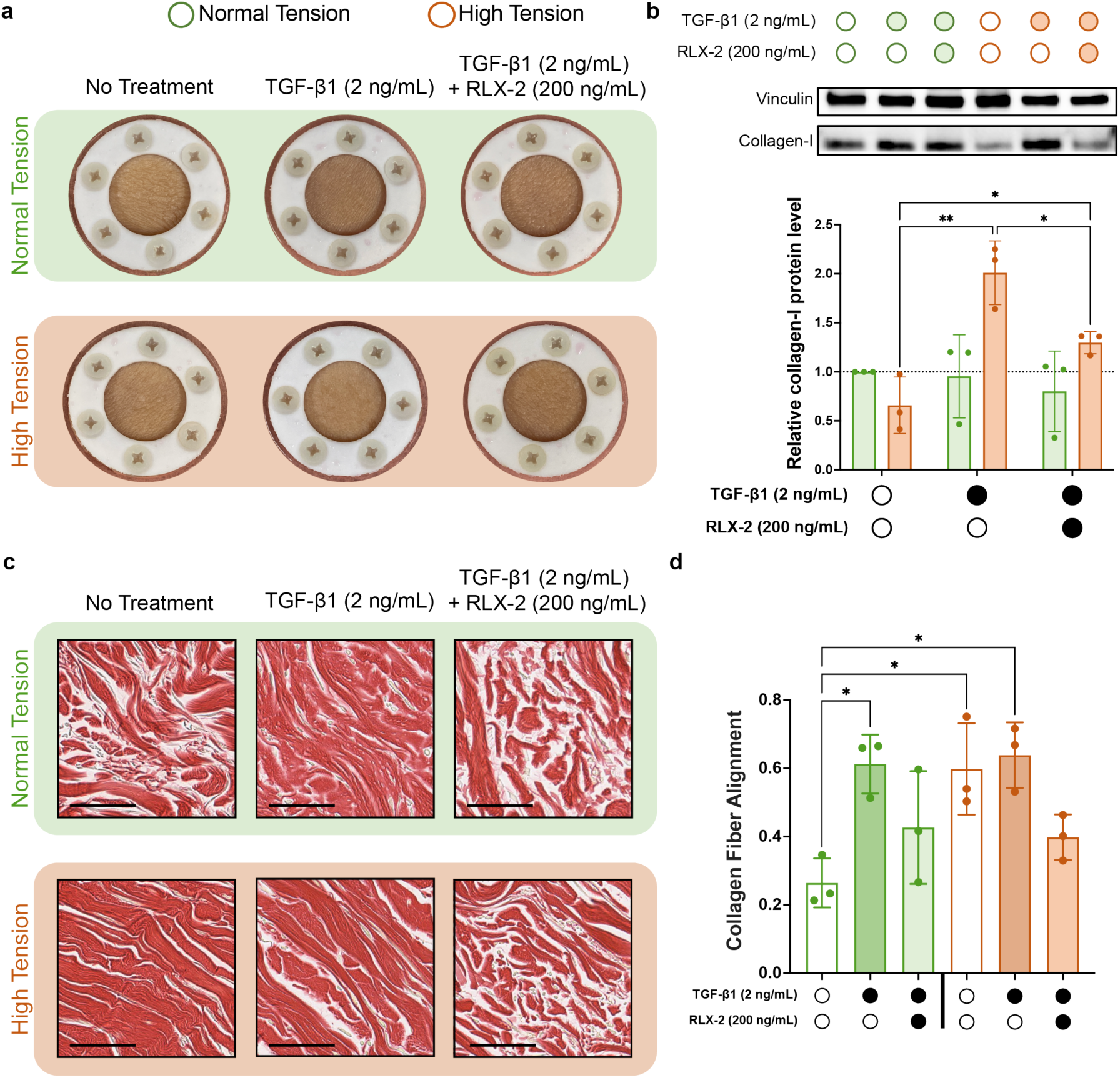
Use of the TenSkin^TM^ ex vivo model to assess the effects of high tension on fibrotic scar development and RLX-2 antifibrotic efficacy. (**A**) Representative images of the TenSkin^TM^ tissue devices under normal physiological and high tension treated *ex vivo* with TGF-β1 (2 ng/mL) with and without RLX-2 (200 ng/mL) for two weeks. (**B**) (Top) Representative western blot of collagen-I from tissues. (Bottom) Densitometry quantification of collagen-I from western blots performed on tissues from three different patients (N=3). (**C**) Representative images of Sirius red stained tissue sections. Scale bar = 50 μm. (**D**) CurveAlign analysis on collagen fibers from Sirius red stained tissue sections. Points represent average of analysis performed on 6 ROIs per tissue section. Bars show standard deviation of averages from 3 different patient samples/experiments (n=6 ROIs, N=3 patients). Bars/circles outlined in green indicate skins under normal tension and bars outlined in orange indicate skins under high tension. Error bars show standard deviation of biological triplicates. Statistical significance was determined with an ordinary one-way ANOVA using a Tukey test to control for multiple comparisons. *=p<0.05, **=p<0.005.

We obtained TenSkin^TM^ from three different patients and maintained half of the samples from each patient under “normal” (physiological) tension and the other half under “high” (pathologic) tension (**Figure 3A**). Under normal tension, TGF-β1 modestly increases collagen-I, and RLX-2 only slightly inhibits collagen-I upregulation. In tissue maintained under increased tension, TGF-β1 stimulation pronouncedly upregulates collagen-I, while RLX-2 markedly suppresses this response, restoring collagen-I levels to those observed under physiological tension. This attenuation of mechanical stress-induced fibrosis by RLX-2 is consistent across all three patient samples (**Figure 3B, S4A**).

Histological analyses further demonstrate that exposure to TGF-β1 and elevated tension induces characteristic pro-fibrotic changes, including increased collagen network alignment and density associated with longer, straighter, and more dense individual fibrils (**Figure 3C-D, S4B**). RLX-2 treatment inhibits these changes to the dermal architecture induced by TGF-β1 at both normal and high tension, promoting maintenance of a normal collagen matrix morphology in the dermal layer of skin, characterized by low-density and basket weave collagen fibrils, even while under both mechanical and biochemical stimulation of pro-fibrotic conditions (**Figure 3C-D, S4**).

Assessment of MMP-1 activity reveals dynamic fluctuations over time, with RLX-2 enhancing active MMP-1 levels relative to TGF-β1 alone at early and later phases, most prominently in the normal tension groups. These changes align with periods critical for limiting contracture and supporting ECM remodeling. Importantly, RLX-2 does not disrupt the natural cycle of MMP-1 regulation, but augments its activity at key stages, thereby preventing both excessive matrix deposition and contracture (**Figure S5**).

Interestingly, RLX-2 increases the relative amount of endogenous active MMP-1 compared to TGF-β1 alone in the first three days of treatment (**Figure S5C**), which is consistent to the timing of RLX-2 efficacy in the *in vitro* models discussed previously. MMP-1 upregulation is also typical of early stages of wound healing during keratinocyte migration to close the wound^41^. We then observed reduced MMP-1 with RLX-2 treatment, which aligns with the re-epithelial stage. The relative MMP-1 ratio increases again, corresponding with ECM remodeling (**Figure S5C**).

### RLX-2 mitigates fibrotic scarring in vivo following burn injury

Wound healing is a complex, multicellular, three-dimensional process with critical interplay between the immune system, fibroblasts, and ECM. To investigate the *in vivo* efficacy of RLX-2 at promoting proper wound healing and ultimately reducing fibrotic scar formation, we used an established *in vivo* murine hypertrophic scar model (**Figure 4A**)^42^. In this chemical burn wound model, an identifiable fibrotic scar is present by day 28. It possesses a thickened epidermis, increased collagen content and alignment in the dermis, and a loss of normal dermal structures such as hair follicles compared to normal mouse skin (**Figure S6**). Together, these scar features resemble the collagen deposition and architecture of a human hypertrophic scar (**Figure S6C**). However, we acknowledge that wound healing in mice differs from that in humans, primarily because tissue contracture plays a greater role than matrix proliferation^43,44^. While murine wound healing and fibrosis models have limitations, they remain valuable for evaluating new anti-fibrotic therapeutic strategies.

**Figure 4.**
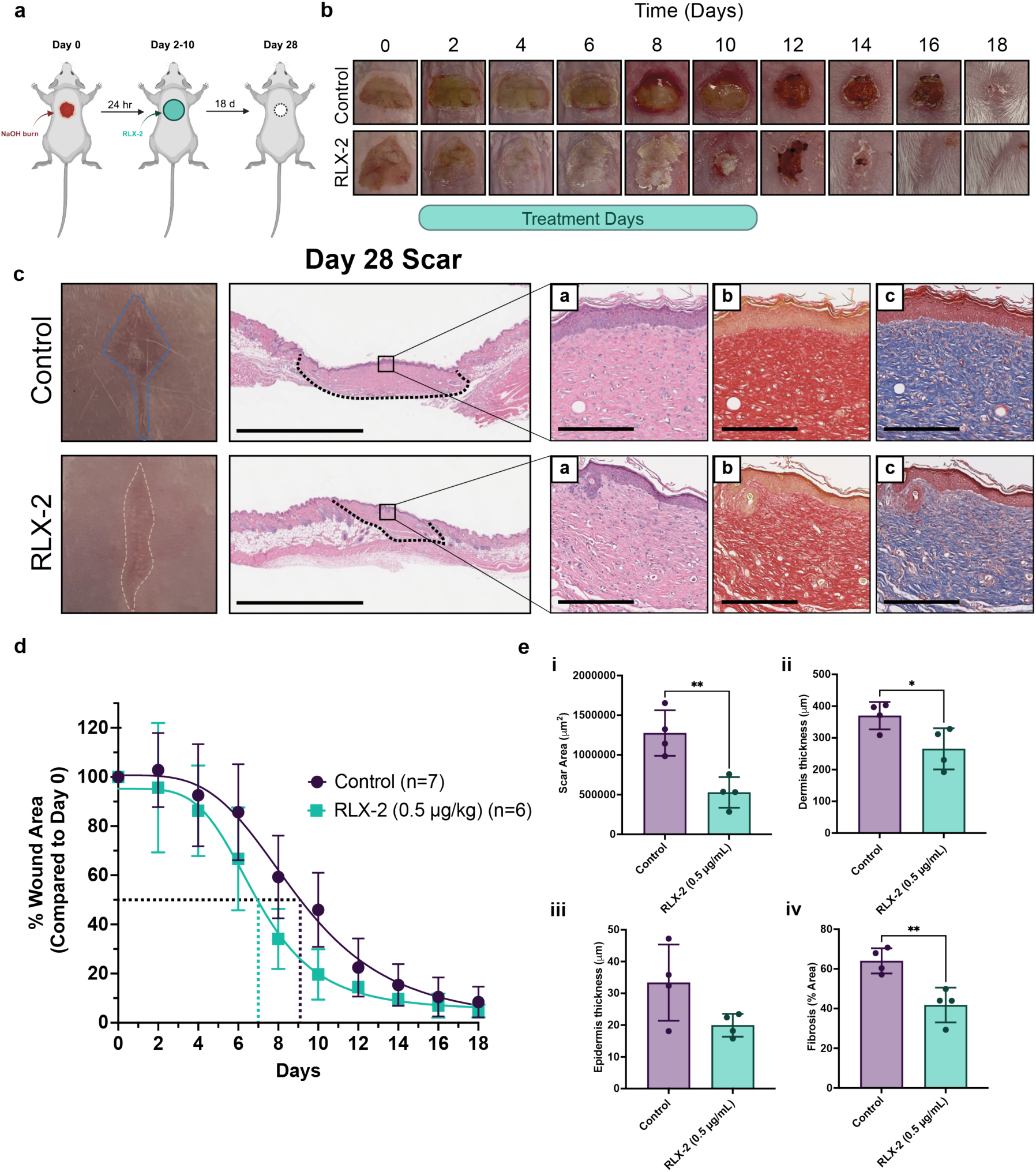
Healing of murine chemical burn wound with multiple treatments of RLX-2. (**A**) Schematic of chemical burn induction, RLX-2 treatment, and wound monitoring over 28-day study (**B**) Representative images of NaOH burn wounds healing over 18 days. Wounds received Whatman paper soaked in 1X PBS (Control) or RLX-2 (0.5 μg/kg, ∼25 – 30 ng RLX-2 per wound) every other day for the first 10 days of healing, beginning at Day 0. (**C**) Representative images of scars at day 28 from (top) control and (bottom) RLX-2-treated wounds with corresponding H&E staining. Dotted lines indicate scar area. Scale bar = 2 mm. Magnified images of the areas in the boxes are shown with (a) H&E, (b) Sirius red, (c) Trichrome staining. Scale bar = 200 μm. (**D**) Quantification of wound area compared to the initial wound area on Day 0. Dotted lines show the day at which each group reached 50% healed. Control (n = 7 wounds): 9.065 days, RLX-2 (0.5 μg/kg) (n=6 wounds): 6.949 days. (**E**) Quantification of (i) scar area from H&E stained tissues sections, (ii) dermis thickness, (iii) epidermis thickness, and (iv) fibrosis percentage from Trichrome stained tissue sections (n=4 wounds).

RLX-2 administration during the initial wound healing phase reduces scar area and accelerates wound closure (**Figure 4B**). RLX-2 treated sites reach 50% healed size 2 days earlier and exhibit lighter scar coloration in comparison with the PBS treated control (**Figure C-D**). Histologically, the RLX-2 treated group shows markedly decreased collagen deposition, a thinner epidermal and dermal layers, and smaller overall scar area compared to controls at day 28 (**Figure 4C, E**).

### RLX-2 decreases collagen density and alignment in ex vivo human hypertrophic scar

To directly address whether RLX-2 modifies established scar architecture—rather than solely prevent fibrotic changes—we examined its effects on fully formed human hypertrophic scars *ex vivo*. Unlike previous experiments that assessed RLX-2’s capacity to prevent or mitigate fibrosis during active wound healing or fibrotic induction, here we cultured biopsy punches obtained from mature hypertrophic scar tissue from a patient and initiated RLX-2 treatment after fibrosis was established. Notably, RLX-2 administered to this preformed scar tissue over a 14-day period significantly reduces collagen fiber alignment and density (**Fig. 5A, C-D**). These alterations are most evident in the lower reticular dermis, as confirmed by a blinded pathologist using a semi-quantitative scoring system (**Fig. 5B, Table S1-4**). These findings demonstrate that RLX-2 can remodel existing fibrotic scar architecture in human skin, highlighting its therapeutic potential not only for scar prevention but also for treating established dermal fibrosis.

**Figure 5.**
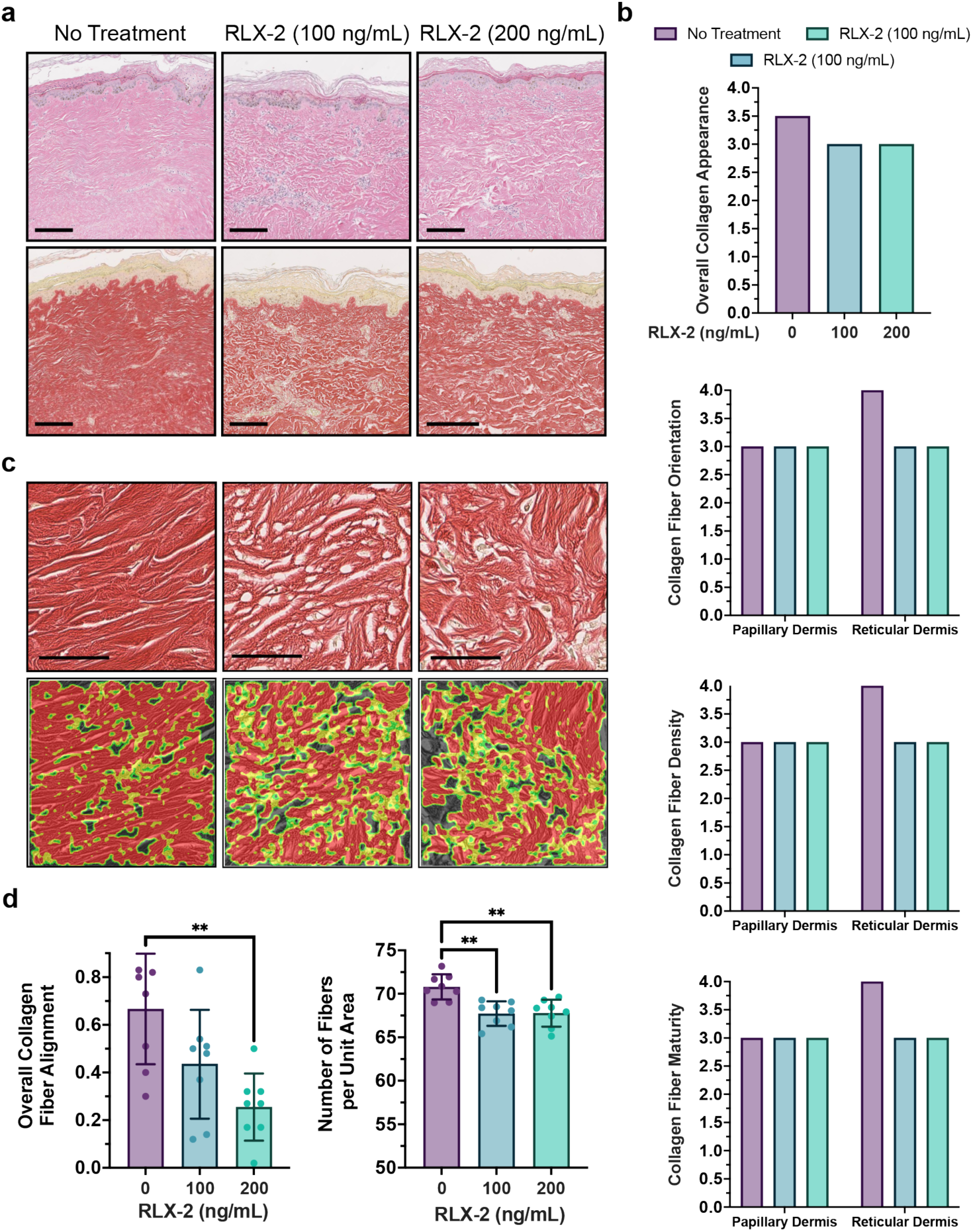
Histological evaluation of hypertrophic scar tissue cultured ex vivo with RLX-2 for 14 days. (**A**) (Top) H&E and (Bottom) Sirius red staining of human hypertrophic scar tissue after *ex vivo* culture with and without RLX-2 (100 or 200 ng/mL) for 14 days. Scale bar = 200 μm (**B**) Pathologist evaluation of collagen fibers from H&E sections. Scores ranged from 0 – 5, with 0 being closest to normal skin and 5 being fibrotic and similar to a keloid scar. Tables 3.1 – 3.4 show assessment criteria used. (**C**) Representative ROIs from Sirius red stained tissue sections with corresponding fiber alignment map from CurveAlign analysis. Red regions on alignment maps indicate higher degree of alignment. Scale bar = 50 μm. (**D**) CurveAlign analyses of overall collagen fiber (Right) alignment and (Left) density. Error bars show standard deviation of 8 ROIs from a single Sirius red tissue section (n=8 ROIs, N=1 patient). Statistical significance was determined by an ordinary one-way ANOVA using a Tukey test to control for multiple comparisons. **=p<0.005.

## Discussion

Fibrotic scarring results from a dysregulated wound healing process, characterized by persistent TGF-β1 signaling, excessive myofibroblast differentiation, and aberrant ECM deposition. In hypertrophic scars specifically, elevated TGF-β1 drives overproduction of collagen-I, collagen-III, and αSMA, while suppressing MMP-1 and MMP-9, leading to a dense, crosslinked collagen fiber network. Inspired by both maternal cervical and uterine remodeling during pregnancy and scarless wound healing observed in utero, we identified endogenous relaxin-2 (RLX-2) as a promising “scarless” therapeutic candidate in both the prevention and treatment of pathologic cutaneous scarring. Across *in vitro*, *ex vivo*, and *in vivo* models, RLX-2 consistently promotes healthy skin architecture, even under pro-fibrotic conditions.

In 2D *in vitro* studies, RLX-2 inhibits TGF-β1-mediated fibroblast-to-myofibroblast differentiation by blocking the Smad2 pathway, normalizing ECM gene and protein expression (**Figure 1**). In a 3D *in vitro* collagen gel model, RLX-2 slows gel contraction at early time points and abrogates excessive ECM production, yet does not fully inhibit contractility, indicating its potential to attenuate fibrosis while preserving effective wound closure and tissue regeneration. *Ex vivo* human skin biopsy experiments reveal dose-dependent effects: at 100 ng/mL, RLX-2 partially restores collagen expression and MMP-1/9 toward normal levels, while reducing collagen fiber alignment and density. At 200 ng/mL, RLX-2 induces a collagen profile more typical of fetal and early wound healing, including increased collagen-III and fibronectin, with additional upregulation of MMP-1/9, suggesting a shift towards regenerative, scarless healing (**Figure 2, S1-S3**). These results demonstrate that RLX-2 dosage can fine-tune the wound healing phenotype, transitioning from fibrotic to scarless tissue characteristics.

RLX-2 also counteracts tension-mediated fibrosis in *ex vivo* human skin models. Under high-tension and TGF-β1 stimulation, RLX-2 preserves a looser and more randomized dermal architecture, reduces overproduction and alignment of collagen fibers, and dynamically regulates MMP-1 levels at critical stages of repair (**Figure 3, S4, S5**). *In vivo* murine models further support RLX-2’s efficacy, with treated wounds exhibiting faster closure, minimized scar area, and improved tissue quality (**Figure 4, S5**). With human translation in mind, preliminary studies conducted on human hypertrophic scar tissue reinforce RLX-2’s capacity to not only prevent but also reverse fibrosis while still permitting normal wound maturation and remodeling (**Figure 5**). These results position RLX-2 as a transformative therapeutic candidate, with the potential to shift clinical paradigms from scar management to scar prevention and skin regeneration.

## Supporting information

suoolemental information

## Acknowledgements

Funding for this work is in part from the NIH (1R01AR083161 – MWG, DR, TPS; T32EB006359 – JRK) and the William Fairfield Warren Professorship at Boston University (MWG). This work was additionally supported by Boston University Micro and Nano Imaging Facility through NIH S10OD024993. We also would like to thank the Foundation for Research on Relaxin in Cardiovascular and other Diseases (RRCA) for the gift of Relaxin-2 and Drs. Ara Nazarian and Edward Rodriguez for helpful comments.

## Data Availability Statement

The data that supports the findings of this study are available upon request.

## CREDIT Authorship Contribution Statement

Conceptualization: AKW, KMH, JRK

Data collection & analysis: AKW, KMH, JRK, RMN

Writing – Original Draft: AKW, KMH, MWG

Writing – Review & Editing: All authors

Funding: TPS, DSR, MWG

Supervision: TPS, DSR, MWG

## Materials and Methods

All reagents were purchased from Thermo-Fisher unless stated otherwise.

### In vitro primary cell culture

All cell culture experiments were performed in a humidified incubator at 37°C and 5% CO2. Primary normal human dermal fibroblasts were acquired from two sources: 1) Commercially purchased Primary Dermal Fibroblast; Normal, Human, Adult (HDFa) (ATCC, PCS-201-012) and cultured in Fibroblast Basal Medium (ATCC, PCS-201-030) with addition of a Low-Serum Fibroblast Growth Kit (2% FBS, ATCC, PCS-201-041) and 1% Penicillin-Streptomycin (PS) (Gibco); 2) isolated from normal skin of patient donors and generously provided by Dr. Andreea Bujor (BUMC). Isolated primary human dermal fibroblasts were cultured in DMEM (Corning, 10013CV) + 10% FBS (R&D Systems) + 1% PS. All experiments were performed between passage 2-10. For western blot and RT-qPCR experiments, cells were plated at 100,00 – 200,000 cells/well in a tissue culture-treated 6-well plate, depending on the length of the treatment. Cells were allowed to adhere overnight and treated at ∼80-90% confluence in low-serum media (0.4% FBS for ATCC normal dermal fibroblasts, 2% FBS for all other cell lines) + 1% PS. For immunostaining experiments, cells were either plated at 100,000 – 200,000 cells/well in a tissue culture-treated 6-well plate containing glass cover slips or at 30,000 cells/well into a 24-well tissue culture-treated plate with a single glass cover slip per well.

### Recombinant human relaxin-2 and TGF-β1

Recombinant human relaxin-2 used in all *in vitro*, *ex vivo* and *in vivo* experiments was obtained as a generous gift from the Relaxin RRCA Foundation (Italy) and was previously validated for activity, purity and structure using a human RLX-2 ELISA, a cAMP activity assay and circular dichroism^19^. EC50 values corresponded to the range with previously published values ^21,45–47^, confirming biologic activity. The RLX-2 used in these studies was also previously validated for purity and size using SDS-PAGE and MALDI-TOF Mass Spectrometry ^17^. RLX-2 was resuspended to 200 μg/mL in sterile 1X PBS, aliquoted and stored at -80°C. TGF-β1 was purchased from Abcam (ab50036) and resuspended in 10mM citric acid, pH 3.0 at 50 μg/mL before diluting to 2 or 1 μg/mL in sterile 1X PBS containing 2 mg/mL BSA. Diluted TGF-β1 was aliquoted and stored at -20°C. All aliquots were either thawed at the time of use or stored at 4°C and used within 1 week. Aliquots were not subjected to additional freeze-thaw cycles. Dilutions for experiments were prepared directly in cell culture media.

### cAMP Assay

Primary human dermal fibroblasts were plated at 12,000 cells/well in a tissue culture treated 96-well white opaque plate and allowed to adhere overnight. The Promega cAMP Glo-Max assay (V1681) was used following the manufacturer protocols. RLX-2 was diluted to 13.5 μg/mL in media and a 3-fold serial dilution was performed to generate the activity curve. Cells were treated with RLX-2, forskolin (10 μM)(Sigma-Aldrich, F3917) or RLX-2 + Forskolin (10 μM) for 40 minutes before measuring cAMP concentration.

### Western blot

2D cells were collected in ice cold 1X PBS and pelleted at 4°C. Supernatant was removed and the pellet was lysed in RIPA buffer containing 1X protease/phosphatase inhibitor cocktail + 1X EDTA. Samples were sonicated briefly using a probe sonicator and centrifuged at 15,000 x G to pellet cell debris. The supernatant was removed and its protein concentration was determined using a Pierce BCA Protein Assay following the manufacturer’s protocol. All samples were diluted to the same protein concentration and volume using RIPA + 1X protease/phosphatase inhibitor cocktail + 1X EDTA. 4X Laemmli sample buffer (Bio-Rad) was added to samples to a final 1X concentration and samples were heated at 95°C for 10 minutes. 1-10 μg of protein/well was loaded onto a Bio-Rad Mini-PROTEAN TGX gel and SDS-PAGE was performed in Tris-Glycine-SDS running buffer. Proteins were transferred to a PVDF membrane (Bio-Rad) using Tris-Glycine buffer + 20% methanol. Membranes were blocked in 5% non-fat dried milk in TBS-T at room temperature for 1 hour. Primary antibodies were diluted in 5% BSA in TBS-T at 1:1000 dilution, except fibronectin, which was diluted 1:500. Membranes were incubated with primary antibodies overnight at 4°C. All washes were performed in TBS-T. Secondary antibodies were diluted at 1:2000 in 5% non-fat dried milk in TBS-T and incubated with membranes for 1 hour at room temperature. Chemiluminescence was read using SuperSignal West Femto or West Atto Maximum Sensitivity substrate on a ChemiDoc XRS+ (Bio-Rad). Primary antibodies: Collagen-I (Novus Biologicals, NB600-408), pro-collagen1α1 (R&D Systems, AF6220), collagen-III (Novus Biologicals, NB600-594), αSMA (Abcam, ab7817), Smad2/3 (Cell Signaling Technology, 8685), P-Smad2 (Cell Signaling Technology, 3108), Cyclophilin-B (Cell Signaling Technology, 43603S), Vinculin (Cell Signaling Technology, 13901), Fibronectin (Santa Cruz Biotechnology, sc-59826). Secondary antibodies: anti-mouse IgG-HRP (Cell Signaling Technology, 7076), anti-rabbit IgG-HRP (Cell Signaling Technology, 7074), anti-sheep IgG-HRP (R&D Systems, HAF016).

### Immunofluorescence staining

Cells were plated at 100,000 – 200,000 cells/well onto glass coverslips placed in a 6-well plate or 30,000 cells/well onto a single cover slip placed in a well of a 24-well plate. Following treatments, cells were rinsed with 1X PBS and fixed in 4% PFA for 15 minutes at room temperature. For cells stained with αSMA and α-actin, an additional permeation step was performed using ice cold methanol at -20°C for 10 minutes. All washes were performed using 1X PBS. Cover slips were blocked in 5% BSA in 1X PBS + 0.3% TritonX-100 for 1 hour at room temperature. All antibodies were diluted in 1% BSA in 1X PBS + 0.3% TritonX-100. Cover slips were incubated in primary antibodies overnight at 4°C. Primary antibodies and dilutions: Collagen-I (Novus Biologicals, NB600-408): 1:200, Smad2/3 (Cell Signaling Technology, 8685): 1:100, αSMA (Abcam, ab7817): 1:100 – 1:500, α-actin (Cell Signaling Technology, 4970): 1:50. Cover slips were incubated with fluorescent AlexaFluor IgG secondary antibodies (Jackson ImmunoResearch) diluted to 1:400 or 1:500 in antibody dilution buffer for 1 hour at room temperature in the dark. F-actin was stained using Phalloidin CruzFluor 633 (Santa Cruz, sc-363796) at 1:1000 dilution in 1X PBS for 10 minutes at room temperature in the dark following secondary antibody incubation and wash steps. Nuclei were stained using Hoechst 33443 dye diluted to 1 μg/mL in 1X PBS and incubated with cells for 10 minutes at room temperature in the dark. Cover slips were washed with 1X PBS following all staining, mounted onto microscope slides using Prolong Gold or Diamond Antifade Mounting Media, and allowed to cure overnight at room temperature in the dark. Additional slides were stained following the same protocol using Mouse IgG1 Isotype Control (R&D Systems, MAB 0002) or Rabbit IgG Isotype Control (Thermo-Fisher, 02-6102) diluted at matched concentrations to the primary antibodies with the corresponding animal source to confirm specific staining. Slides were imaged using an Olympus FV1000 or FV3000 Scanning Confocal Microscope in the Micro/Nano Imaging (MNI) Facility at Boston University. For quantifications of Smad2/3 nuclear localization, Smad2/3 fluorescence signal was scored as primarily nuclear versus evenly distributed/predominantly cytoplasmic in 100-200 cells for each experimental condition, following methods in ^48^.

### RNA extraction and RT-qPCR

For *in vitro* cell culture experiments, mRNA extraction was performed using the Qiagen RNeasy Plus Kit (74134) following the manufacturer protocols. Following extraction, RNA concentration and purity were assessed by the A260/A280 values using a Nanodrop 2000 (Thermo-Fisher). Samples were normalized to the same concentration of RNA and cDNA was generated using the High-Capacity cDNA Reverse Transcription Kit (Applied Biosystems, 4368814) following the manufacturer’s protocol with a Bio-Rad T100 Thermal Cycler. qPCR was performed using TaqMan Fast Advanced Master Mix (Applied Biosystems, 4444556) following the manufacturer’s protocol and TaqMan probes for specified genes. qPCR was run on an Applied Biosystems StepOne Plus RT-PCR Thermal Cycler in the Bio-Interface Technologies (BIT) facility at Boston University. TaqMan probes (Applied Biosystems): Collagen-I (Hs00164004_m1), Collagen-III (Hs00943809_m1), αSMA (Hs00426835_g1), CTGF (Hs00170014_m1), MMP-1 (Hs00899658_m1), MMP-9 (Hs00957562_m1), TIMP-1 (Hs01092512_g1), GAPDH (Hs02786624_g1), B2M (Hs99999907_m1). All probes were FAM-MGB or VIC-MGB. In some experiments, two probes with different fluorophores were multiplexed to measure multiple genes in a single well. Relative gene expression was determined by calculating the ΔΔCT values using the corresponding endogenous control reference gene (GAPDH or B2M) and the untreated control group in each experiment. Efficiency was assumed to be 2 for the 2^-ΔΔCT^ calculation. Statistical analysis was performed using GraphPad Prism 10. For comparison between two groups, statistical significance was calculated using a two-tailed unpaired T-test. For comparison between 3 or more groups, statistical significance was determined by an ordinary one-way ANOVA using a Tukey test to control for multiple comparisons. Data is reported as mean + standard deviation with significance reported for adjusted p<0.05. *=p<0.05, **=p<0.005, ***=p<0.0005, ****=p<0.0001

### Fibroblast-embedded collagen gels

All collagen gels were made from rat tail collagen-I at a final concentration of 3 mg/mL. Collagen gels were prepared using two methods based on the source of rat-tail collagen-I: 1) High concentration rat-tail collagen-I (Corning, 354249) was diluted with 0.2N acetic acid and mixed with 2X PBS + 100 mM HEPES as a neutralization buffer or 2) RatCol Rat Tail Collagen (Advanced Biomaterials, 5153) was mixed with the corresponding neutralization buffer (Advanced Biomaterials) in a 9:1 ratio and kept on ice. Cells were trypsinized and resuspended in complete media to a concentration allowing for 300,000 – 500,000 cells/gel. The cell suspension was added to the collagen mixture and pipetted gently and thoroughly. 2.5 mL/well of the collagen-cell suspension was added to a 3.0 μm pore PET or Polycarbonate transwell insert (Corning, 353091/Corning 3414) placed in either a regular tissue culture 6-well plate or a deep-well 6-well plate (Fisher, 08-774-183). Gels were placed in a humidified incubator at 37°C and 5% CO2 for 1-3 hours for collagen polymerization. Once gels appeared opaque, complete media containing the indicated treatments was added to the top and bottom of the insert so that the entire gel was submerged. Media and treatments were changed every 2-3 days for the duration of the experiment. Pictures were taken of gels at each media change and diameters and areas were measured using ImageJ software based on the dimensions of the transwell. Percent contraction was calculated as the difference between the diameter of the transwell insert and the measured diameter of the gel, divided by the diameter of the insert, multiplied by 100. 0% contraction indicates the gel is the same size as the transwell insert and no contraction occurred. 50% contraction indicates the gel contracted to half the original diameter of the transwell insert. To combine multiple experimental replicates, the percent contraction of gels was graphed versus the number of days after an initial change in gel size was observed. The first measurement of contraction was Day 1 post start of gel contraction. Rate of contraction was calculated as the change in diameter of the collagen gels divided by the days in between measurement times. Statistics were calculated and regression analysis was performed using GraphPad Prism 10, where applicable.

### Protein isolation from fibroblasts in collagen gels

Collagen gels were submerged in RIPA buffer containing 1X protease/phosphatase inhibitor cocktail + 1X EDTA. The gels were lysed by alternating between vortexing, sonicating for 5 – 10 seconds at 20% amplitude with a probe sonicator, and incubating on ice until the gel was broken up into a slurry. The mixture was centrifuged at 10,000 x G for 10 minutes at 4°C to pellet the remaining collagen debris. The supernatant was removed and used for western blot analysis. Protein quantification, sample preparation and western blot were carried out same as previously described.

### Ex Vivo culture of human skin tissues

Human normal skin or hypertrophic scar tissue was excised from patients receiving abdominoplasty surgery and collected as discarded tissue by Dr. Daniel Roh at Boston University Chobanian & Avedisian School of Medicine in accordance with Boston University Institutional Review Board (IRB) protocol numbers H-39933 and H-41935. All experiments involving biological specimens were approved by the Institutional Biosafety Committee (IBC) at Boston University. Researchers acquiring tissue samples were blinded to all identifying patient information. The hypertrophic scar sample was also removed during an abdominoplasty. All tissue handling and *ex vivo* culture was performed under sterile aseptic conditions in a biosafety cabinet.

#### Normal skin tissue preparation and culture

Tissue was washed 3X in sterile 1X PBS. Subcutaneous fat was removed, and tissue was submerged in DMEM + 10% FBS + 1% PS for 30 minutes at 37°C. 8 mm biopsy punches were cut out of the skin samples. The hypertrophic scar was small, so 4 mm biopsy punches were made along the scar. Rat tail collagen-I was mixed with neutralization buffer (Advanced Biomaterials, 5153) in a 9:1 ratio to a final collagen concentration of 2 mg/mL. For normal skin tissues, 500 μL of the neutralized collagen solution was added to a transwell insert in a 6-well cell culture plate (24 mm diameter, 3 μm pore, polycarbonate, ThermoFisher, 140642). For hypertrophic scar punches, 100 μL of neutralized collagen solution was added to a transwell insert in a 24-well plate (3 μm pore, polycarbonate, ThermoFisher, 141004). Each biopsy punch was placed in a separate transwell insert in the collagen solution with the epidermis facing upwards. The plates were placed in a 37°C humidified incubator with 5% CO2 for 15-20 minutes to allow the collagen to polymerize. Media: DMEM + 10 % FBS + 1% PS + hydrocortisone (Lonza, cc-4031G), gentamicin sulfate + amphotericin-B (Lonza, GA-1000) + 5 μg/mL insulin. For normal skin punches, 2 – 3 mL media was added to the outside of the transwell insert and 100 μL was added inside the insert above the collagen layer and surrounding the tissue. For hypertrophic scar tissue, 1.5 mL media was added below the transwell and 50 μL was added around the tissue. The epidermis was not submerged in media to maintain an air-liquid interface. Tissues were allowed to rest overnight in a 37°C humidified incubator with 5% CO2. The next day, treatments were diluted directly in the media. Media was removed from the plate, collected, and media containing treatments were added with 2 mL outside the transwell insert and 200 μL inside for normal skin punches (2.2 mL total) and 1.5 mL outside and 50 μL inside for hypertrophic scar punches (1.55 mL total). Every 2 days, 600 μL – 1 mL media was collected and stored at -80°C for future analyses. The remaining media was removed from the wells and fresh media containing treatments was added. Tissues were treated every other day for 14 days.

#### TenSkin^TM^ Tissues

Skin tissues maintained under tension were commercially purchased from TenBio (ten-bio.com). Skin tissues were harvested from abdominoplasty procedures. Three different donors were used. Six TenSkin^TM^ models were generated from each donor. Three were put under “normal” physiological tension and three were put under “high” tension. The models were shipped fully prepared from TenBio in a media gel. Upon receiving the models, all steps were followed according to the company protocols. The models were removed from the gels and placed in a 60 mm cell culture dish with 7.5 mL of media provided by TenBio, ensuring no bubbles were trapped underneath. The models were acclimated in a 37°C, 5% CO2 incubator for 30 minutes. TGF-β1 and/or RLX-2 were added directly to the media in the dishes for a final concentration of 2 ng/mL and 200 ng/mL, respectively. The models were moved to a new 60 mm dish containing fresh media with treatments every 24 hours for 2 weeks. A 1.8 mL sample of media was collected each day and stored at -80°C for additional analyses.

### MMP-1 Activity Assay

Media samples were thawed on ice and centrifuged at 10,000 x G for 15 minutes at 4°C to remove any particulates. Supernatants were transferred to new tubes and used for the assay. MMP-1 concentration was quantified using the Fluorokine E Human Active MMP-1 Fluorescent Activity Assay (R&D, F1M00), following the manufacturer protocol. p-Aminophenylmercuric Acetate (APMA) was not added to the samples in order to measure endogenous levels of active MMP-1 in media samples. Fluorescence was measured at λEx = 320 nm/λEm = 405 nm on a Spectramax iD3 plate reader (Molecular Devices). The standard curve was used to calculate MMP-1 concentration in media samples.

### Ex vivo tissue collection

Media was removed from wells, tissues were rinsed in ice cold 1X PBS, and excess collagen was removed. Biopsy punches were weighed and either placed directly into 10% phosphate buffered formalin for histological assessment or cut into 3-4 pieces depending on the analysis being performed. Each tissue piece was weighed before storing. For pieces being used for RT-qPCR, tissues were submerged in RNAprotect Tissue Reagent (Qiagen, 76104) at a volume 10X the tissue mas (e.g., 30 mg tissue = 300 μL reagent). Tissues in RNAprotect were stored at 4°C for at least 24 hours before storing at -20°C or removing the reagent and storing the preserved tissues at -80°C. For tissues being processed for protein extraction, western blot or collagen analysis, pieces were flash frozen in liquid nitrogen and stored at -80°C. In experiments with hypertrophic scars, whole biopsy punches were used for each analysis method. For the TenSkin^TM^ models, 4 mm or 6 mm biopsy punches were made throughout the tissue in the center of the device. Each biopsy punch was stored for either histology, protein, RNA, or collagen quantification using the same methods as described above.

### RNA extraction from skin tissues

Tissues were thawed, cut and weighed into pieces under 30 mg. Tissue piece was placed in a screw cap tube with either a 5 mm stainless steel bead (Qiagen, 69989) or prefilled 2.0 mL tubes with garnet shards and a 6 mm zirconium bead (Benchmark Scientific, D1033-30G). 300 μL of RLT Buffer containing β-mercaptoethanol was added to each tube. Tubes were placed in a Bead Bug 6 Homogenizer (Benchmark Scientific) and homogenized at a speed of 4000 for one minute at a time, alternating with incubation on ice until tissue pieces were homogenized. Tubes were centrifuged briefly to remove beads and the supernatant was centrifuged through Qiashredder columns (Qiagen, 79656) to further homogenize the lysates. RNA extraction was performed using a Fibrous Tissue RNA Extraction Kit (Qiagen, 74704) following the manufacturer protocol. RNA quality and concentration were assessed by the A260/A280 ratio measured using a Nanodrop2000 (ThermoFisher). If the A260/A280 ratio was low, RNA was cleaned and concentrated using the RNeasy MiniElute Cleanup Kit following the manufacturer protocol. Reverse transcription and qPCR were performed as previously described.

### Protein extraction from skin tissues

Tissues were kept on dry ice and a 5-10 mg piece of tissue was cut into smaller pieces using a scalpel. Tissues were either homogenized using a Dounce homogenizer on ice or a Bead Bug 6 Homogenizer (Benchmark Scientific) with a 5 mm stainless steel bead (Qiagen, 69989) or prefilled 2.0 mL tubes with garnet shards and a 6 mm zirconium bead (Benchmark Scientific, D1033-30G). Tissues were lysed in ice-cold RIPA buffer with 1X protease/phosphatase inhibitor cocktail + 1X EDTA at 300 µL/5 mg tissue. For tissues homogenized with the Bead Bug, tissue lysates were incubated on an orbital shaker at 4°C for 2 hours. Lysates were transferred to new tubes and sonicated using a probe sonicator at 20% amplitude for 5-10 seconds at a time, alternating with incubations on ice. Lysates were centrifuged at 15,000 x G for 15 minutes at 4°C to remove debris and insoluble collagen. Supernatants were removed and used for Western blots. Western blots were performed according to the procedure previously described. Vinculin was used as the endogenous loading control protein for skin tissue western blots.

### Collagen Assays

For soluble collagen content in the media, samples were thawed and centrifuged at 10,000 x G for 15 minutes at 4°C. Supernatants were transferred to new tubes and analyzed using a Soluble Collagen Assay (Sigma-Aldrich, CS006) following the manufacturer’s protocol. Fluorescence was measured at λEx = 375 nm/λEm = 465 nm on a Spectramax iD3 plate reader (Molecular Devices). A hydroxyproline assay (Sigma-Aldrich, MAK357) was used to measure the total collagen content in the tissue samples following the manufacturer protocol. Tissues were kept on dry ice while cutting ∼10 mg for the assay. A probe sonicator was used at 20% amplitude for the initial homogenization step. Samples were measured in duplicate on the assay plate. The final absorbance reading was measured at 560 nm on a Spectramax iD3 plate reader (Molecular Devices).

### In vivo murine model

All experiments involving mice were approved by the Boston University Institutional Animal Care and Use Committee (IACUC) under protocol 201900023. Chemical burn wound procedure was followed based on Kim et al. 2018 ^42^. All experiments were performed using 6-week old female ICR (CD-1) mice (22 – 27g, Charles River Laboratories, Inc.). 8 mm and 10 mm circles were cut from Whatman filter paper (Grade 1, Thermo-Fisher) and sterilized under UV light overnight. A 2N NaOH (Emprove bio, 1370201000) solution was prepared under aseptic conditions using sterile UltraPure Distilled water (Invitrogen). The final solution was filtered through a 0.2 µm syringe filter. Mice were anesthetized with isoflurane using standard procedures and given a subcutaneous injection of 72-hour sustained-release buprenorphine. The hair on the dorsum was shaved and further removed with Nair hair removal cream. To create the wound, an 8 mm Whatman paper circle was fully submerged in the 2N NaOH solution and applied to the back of the mouse for 60 seconds. Two wounds were made per mouse, one on the upper dorsum and one in the center. The initial wound was photographed as the “Day 0” timepoint. The wounds were left open to air overnight. We previously found that RLX-2 requires localized and repeated application for antifibrotic efficacy ^19^. Thus, in this experiment, we applied five doses of RLX-2 (0.5 µg/kg diluted in sterile 1X PBS) every other day for the first 10 days of wound healing. The day after generating the NaOH burn (Day 0), 17 µL of either sterile 1X PBS or the diluted RLX-2 was pipetted onto a 10 mm Whatman paper circle to saturate it. Each 10 mm paper was applied to one wound. The paper circles were covered with a piece of Tegaderm (3M), wrapped with a piece of self-adhesive bandage and secured with medical tape (Johnson & Johnson). Every two days for 10 days, the bandages were removed, the wounds were photographed, and the treatments and bandages were reapplied. All procedures were performed under isoflurane general anesthesia. After 10 days, all bandages were removed and the wounds were allowed to heal open to air until day 28. Wounds were photographed every two days until the hair was completely grown back and they were no longer visible (∼ Day 18). At day 28, the hair was shaved and removed with Nair hair removal cream and the scar was photographed. Mice were euthanized following standard procedures and skin tissue with the scar present was excised and immediately fixed in 10% phosphate buffered formalin for histological analyses.

### Histology

At the conclusion of the *ex vivo* studies, biopsy punches being analyzed for histology were rinsed in 1X PBS and placed immediately in 10% phosphate buffered formalin. Tissues were shipped the same day or the next day to HistoWiz, Inc. for processing and staining. Histology was performed by HistoWiz, Inc. (histowiz.com) using a Standard Operating Procedure and fully automated workflow. Samples were processed, embedded in paraffin, and sectioned at 4μm. HistoWiz, Inc. performed H&E, Masson’s Trichrome and Sirius Red staining of tissue sections. Whole slide scanning (40x) was performed on an Aperio AT2 (Leica Biosystems) and uploaded to the researchers’ personal account on the HistoWiz website.

### Histological analysis

To quantify collagen fiber alignment and density, images of tissue sections were downloaded from histowiz.com and viewed using QuPath to pick regions of interest (ROI) for the analyses. Sirius red stained tissue sections were used for quantification. Each ROI was 512 x 512 pixels in size and imported into the open-source program CurveAlign 4.0 (University of Wisconsin-Madison) ^49–51^. CurveAlign can be used to measure bulk collagen fiber attributes. Measurements of overall alignment and box density were used for analysis. Alignment is calculated based on the length of the sum of orientation vectors divided by the number of absolute angles with respect to the horizontal axis and ranges from 0-1. A value of 1 indicates that all fibers are aligned in a single direction and values closer to 0 indicate lower fiber alignment. The density is calculated based on the number of objects (e.g., fibers) in a square 32 pixels x 32 pixels whose center is an object. It indicates the number of fibers surrounding each other in a defined area. 8 ROIs were analyzed per tissue section (n=8). For normal skin tissues treated with TGF-β1 (1 and 2 ng/mL), ROIs were averaged for each tissue section per patient and graphed together. Two different patient samples were used for normal skin (N=2). Statistical significance was determined in GraphPad Prism 10 by an ordinary one-way ANOVA using a Tukey test to control for multiple comparisons. Data is reported as mean + standard deviation with significance reported for adjusted p<0.05. *=p<0.05, **=p<0.005, ***=p<0.0005, ****=p<0.0001

Pathological scoring of H&E stained tissue sections of hypertrophic scars was performed by Dr. Rosalynn Nazarian, MD (MGH). Dr. Nazarian is a surgical pathologist specializing in fibrotic skin diseases. Scoring criteria are defined in Tables S1-S4 and were adapted from Ozog, D.M., et al., 2013 ^52^, Khatery B.H.M., et al., 2022 ^53^ and Beausang, E., et al., 1998 ^35^. Scores range from 0-5 with 0 indicating collagen fibers resembling those in normal skin tissue and 5 indicating looking fibrotic and keloid-like. For fiber orientation, density and maturity, collagen fibers were assessed separately in the papillary dermis and reticular dermis. Dr. Nazarian was blinded to the treatments applied to the tissues when performing the analysis.

## References

1 Gauglitz, G. G., Korting, H. C., Pavicic, T., Ruzicka, T. & Jeschke, M. G. Hypertrophic scarring and keloids: pathomechanisms and current and emerging treatment strategies. Mol Med 17, 113–125 (2011). 10.2119/molmed.2009.00153

2 Plikus, M. V. et al. Fibroblasts: Origins, definitions, and functions in health and disease. Cell 184, 3852–3872 (2021). 10.1016/j.cell.2021.06.024

3 Younesi, F. S., Miller, A. E., Barker, T. H., Rossi, F. M. V. & Hinz, B. Fibroblast and myofibroblast activation in normal tissue repair and fibrosis. Nat Rev Mol Cell Biol 25, 617–638 (2024). 10.1038/s41580-024-00716-0

4 Lee, W. J., Kim, Y. O., Choi, I. K., Rah, D. K. & Yun, C. O. Adenovirus-relaxin gene therapy for keloids: implication for reversing pathological fibrosis. Br J Dermatol 165, 673–677 (2011). 10.1111/j.1365-2133.2011.10439.x

5 Safonov, I. & Safonov, I. Pathological Scars (Keloid and Hypertrophic Scars). Atlas of Scar Treatment and Correction, 97–160 (2012). 10.1007/978-3-642-29196-8_2

6 Karppinen, S.-M., Heljasvaara, R., Gullberg, D., Tasanen, K. & Pihlajaniemi, T. Toward understanding scarless skin wound healing and pathological scarring. F1000 Research 8, 1–11 (2019).

7 Jeschke, M. G. et al. Scars. Nat Rev Dis Primers 9, 64 (2023). 10.1038/s41572-023-00474-x

8 Ogawa, R. Keloid and Hypertrophic Scars Are the Result of Chronic Inflammation in the Reticular Dermis. International Journal of Molecular Sciences 18, 1–10 (2017). 10.3390/ijms18030606

9 Xue, M. & Jackson, C. J. Extracellular Matrix Reorganization During Wound Healing and Its Impact on Abnormal Scarring. Advances in Wound Care 4, 119–136 (2015). 10.1089/wound.2013.0485

10 Ghazawi, F. M., Zargham, Ramin, Gilardino, Mirko, S., Sasseville, Denis, Jafarian, Fatemeh. Insights into the Pathophysiology of Hypertrophic Scars and Keloids: How Do They Differ? Advances in Skin & Wound Care 31, 582–594 (2017).

11 Mascharak, S. et al. Preventing Engrailed-1 activation in fibroblasts yields wound regeneration without scarring. Science 372 (2021). 10.1126/science.aba2374

12 Reish, R. G. & Eriksson, E. Scars: A Review of Emerging and Currently Available Therapies. Plastic and Reconstructive Surgery 122, 1068–1078 (2008). 10.1097/PRS.0b013e318185d38f

13 Samuel, C. S., Summers, R. J. & Hewitson, T. D. Antifibrotic Actions of Serelaxin – New Roles for an Old Player. Trends in Pharmacological Sciences 37, 485–497 (2016). 10.1016/j.tips.2016.02.007

14 Samuel, C. S. et al. Anti-fibrotic actions of relaxin. Br J Pharmacol 174, 962–976 (2017). 10.1111/bph.13529

15 Chow, B. S. et al. Relaxin signals through a RXFP1-pERK-nNOS-NO-cGMP-dependent pathway to up-regulate matrix metalloproteinases: the additional involvement of iNOS. PLoS One 7, e42714 (2012). 10.1371/journal.pone.0042714

16 Khanna, D. et al. Recombinant human relaxin in the treatment of systemic sclerosis with diffuse cutaneouss involvement: A randomized, double-blind, placebo-controlled trial. Arthritis & Rheumatism 60, 1102–1111 (2009). 10.1002/art.24380

17 Blessing, W. A. et al. Intraarticular injection of relaxin-2 alleviates shoulder arthrofibrosis. Proc Natl Acad Sci U S A 116, 12183–12192 (2019). 10.1073/pnas.1900355116

18 Papworth, M. et al. A novel long-acting relaxin-2 fusion, AZD3427, improves cardiac performance in non-human primates with cardiac dysfunction. Cardiovasc Res 121, 871–881 (2025). 10.1093/cvr/cvaf031

19 Kirsch, J. R. et al. Minimally invasive, sustained-release relaxin-2 microparticles reverse arthrofibrosis. Sci Transl Med 14 (2022). 10.1126/scitranslmed.abo3357

20 Almeida-Pinto, N., Dschietzig, T. B., Brás-Silva, C. & Adão, R. Cardiovascular effects of relaxin-2: therapeutic potential and future perspectives. Clinical Research in Cardiology 113, 1137–1150 (2024). 10.1007/s00392-023-02305-1

21 Erlandson, S. C. et al. The relaxin receptor RXFP1 signals through a mechanism of autoinhibition. Nature Chemical Biology 19, 1013–1021 (2023). 10.1038/s41589-023-01321-6

22 Patil, N. A. et al. Relaxin family peptides: structure-activity relationship studies. British Journal of Pharmacology 174, 950–961 (2017). 10.1111/bph.13684

23 Hossain, M. A. et al. The minimal active structure of human relaxin-2. J Biol Chem 286, 37555–37565 (2011). 10.1074/jbc.M111.282194

24 Bathgate, R. A. D. et al. Relaxin Family Peptides and Their Receptors. Physiol Rev 93, 405–480 (2013). 10.1152/physrev.00001.2012

25 Fink, S. P., Mikkola, D., Willson, J. K. & Markowitz, S. TGF-beta-induced nuclear localization of Smad2 and Smad3 in Smad4 null cancer cell lines. Oncogene 22, 1317–1323 (2003). 10.1038/sj.onc.1206128

26 Grinnell, F. Fibroblast–collagen-matrix contraction: growth-factor signalling and mechanical loading. Trends in Cell Biology 10, 362–365 (2000). 10.1016/s0962-8924(00)01802-x

27 Blokland, K. E. C. et al. Substrate stiffness engineered to replicate disease conditions influence senescence and fibrotic responses in primary lung fibroblasts. Frontiers in Pharmacology 13, 989169 (2022). 10.3389/fphar.2022.989169

28 Discher, D. E., Janmey, P. & Wang, Y. L. Tissue cells feel and respond to the stiffness of their substrate. Science 310, 1139–1143 (2005). 10.1126/science.1116995

29 Wells, R. G. Tissue mechanics and fibrosis. Biochimica et Biophysica Acta (BBA) - Molecular Basis of Disease 1832, 884–890 (2013). 10.1016/j.bbadis.2013.02.007

30 Hinz, B., Mastrangelo, D., Iselin, C. E., Chaponnier, C. & Gabbiani, G. Mechanical tension controls granulation tissue contractile activity and myofibroblast differentiation. Am J Pathol 159, 1009–1020 (2001). 10.1016/S0002-9440(10)61776-2

31 Aarabi, S. et al. Mechanical load initiates hypertrophic scar formation through decreased cellular apoptosis. FASEB J 21, 3250–3261 (2007). 10.1096/fj.07-8218com

32 Zhang, Q. et al. Collagen gel contraction assays: From modelling wound healing to quantifying cellular interactions with three-dimensional extracellular matrices. European Journal of Cell Biology 101, 151253 (2022). 10.1016/j.ejcb.2022.151253

33 Montesano, R. & Orci, L. Transforming growth factor beta stimulates collagen-matrix contraction by fibroblasts: implications for wound healing. Proc Natl Acad Sci U S A 85, 4894–4897 (1988). 10.1073/pnas.85.13.4894

34 Larson, B. J., Longaker, M. T. & Lorenz, H. P. Scarless Fetal Wound Healing: A Basic Science Review. Plastic and Reconstructive Surgery 126, 1172–1180 (2010). 10.1097/PRS.0b013e3181eae781

35 Beausang, E., Floyd, H., Dunn, K. W., Orton, C. I. & Ferguson, M. W. A new quantitative scale for clinical scar assessment. Plast Reconstr Surg 102, 1954–1961 (1998). 10.1097/00006534-199811000-00022

36 Harn, H. I. C. et al. The tension biology of wound healing. Experimental Dermatology 28, 464–471 (2019). 10.1111/exd.13460

37 Junker, J. P. E., Kratz, C., Tollbäck, A. & Kratz, G. Mechanical tension stimulates the transdifferentiation of fibroblasts into myofibroblasts in human burn scars. Burns 34, 942–946 (2008). 10.1016/j.burns.2008.01.010

38 Horowitz, J. C. & Thannickal, V. J. Mechanisms for the Resolution of Organ Fibrosis. Physiology 34, 43–55 (2019). 10.1152/physiol.00033.2018

39 Friedman, S. L., Sheppard, D., Duffield, J. S. & Violette, S. Therapy for fibrotic diseases: Nearing the starting line. Science Translational Medicine 5, 1–17 (2013). 10.1126/scitranslmed.3004700

40 Hinz, B. The extracellular matrix and transforming growth factor-beta1: Tale of a strained relationship. Matrix Biol 47, 54–65 (2015). 10.1016/j.matbio.2015.05.006

41 Caley, M. P., Martins, V. L. & O’Toole, E. A. Metalloproteinases and Wound Healing. Adv Wound Care (New Rochelle) 4, 225–234 (2015). 10.1089/wound.2014.0581

42 Kim, M., Kim, H. & Kang, H. W. Comparative evaluations of hypertrophic scar formation in in vivo models. Lasers Surg Med 50, 661–668 (2018). 10.1002/lsm.22783

43 Li, J. et al. Experimental models for cutaneous hypertrophic scar research. Wound Repair Regen 28, 126–144 (2020). 10.1111/wrr.12760

44 Wong, V. W., Sorkin, M., Glotzbach, J. P., Longaker, M. T. & Gurtner, G. C. Surgical approaches to create murine models of human wound healing. J Biomed Biotechnol 2011, 969618 (2011). 10.1155/2011/969618

45 Illiano, S. et al. Characterization of a new potent and long-lasting single chain peptide agonist of RXFP1 in cells and in vivo translational models. Scientific Reports 12, 1–17 (2022). 10.1038/s41598-022-24716-2

46 Hossain, M. A. et al. A single-chain derivative of the relaxin hormone is a functionally selective agonist of the G protein-coupled receptor, RXFP1. Chemical Science 7, 3805–3819 (2016). 10.1039/c5sc04754d

47 Erlandson, S. C. et al. Engineering and Characterization of a Long-Half-Life Relaxin Receptor RXFP1 Agonist. Molecular Pharmaceutics 21, 4441–4449 (2024). 10.1021/ACS.MOLPHARMACEUT.4C00368

48 Dupont, S. et al. Role of YAP/TAZ in mechanotransduction. Nature 474, 179–183 (2011). 10.1038/nature10137

49 Liu, Y., Keikhosravi, A., Mehta, G. S., Drifka, C. R. & Eliceiri, K. W. Methods for Quantifying Fibrillar Collagen Alignment. Methods in molecular biology (Clifton, N.J.) 1627, 429 (2017). 10.1007/978-1-4939-7113-8_28

50 Bredfeldt, J. S. et al. Automated quantification of aligned collagen for human breast carcinoma prognosis. J Pathol Inform 5, 28 (2014). 10.4103/2153-3539.139707

51 Chen, K. et al. Disrupting mechanotransduction decreases fibrosis and contracture in split-thickness skin grafting. Science Translational Medicine 14, 1–17 (2022). 10.1126/scitranslmed.abj9152

52 Ozog, D. M. et al. Evaluation of clinical results, histological architecture, and collagen expression following treatment of mature burn scars with a fractional carbon dioxide laser. JAMA Dermatol 149, 50–57 (2013). 10.1001/2013.jamadermatol.668

53 Khatery, B. H. M., Hussein, H. A., Abd-El-Raheem, T. A., El Hanbuli, H. M. & Yassen, N. N. Assessment of intralesional injection of botulinum toxin type A in hypertrophic scars and keloids: Clinical and pathological study. Dermatol Ther 35, e15748 (2022). 10.1111/dth.15748

