## Supplementary material for "Relaxin-2 drives regenerative healing and suppresses scar formation": suoolemental information

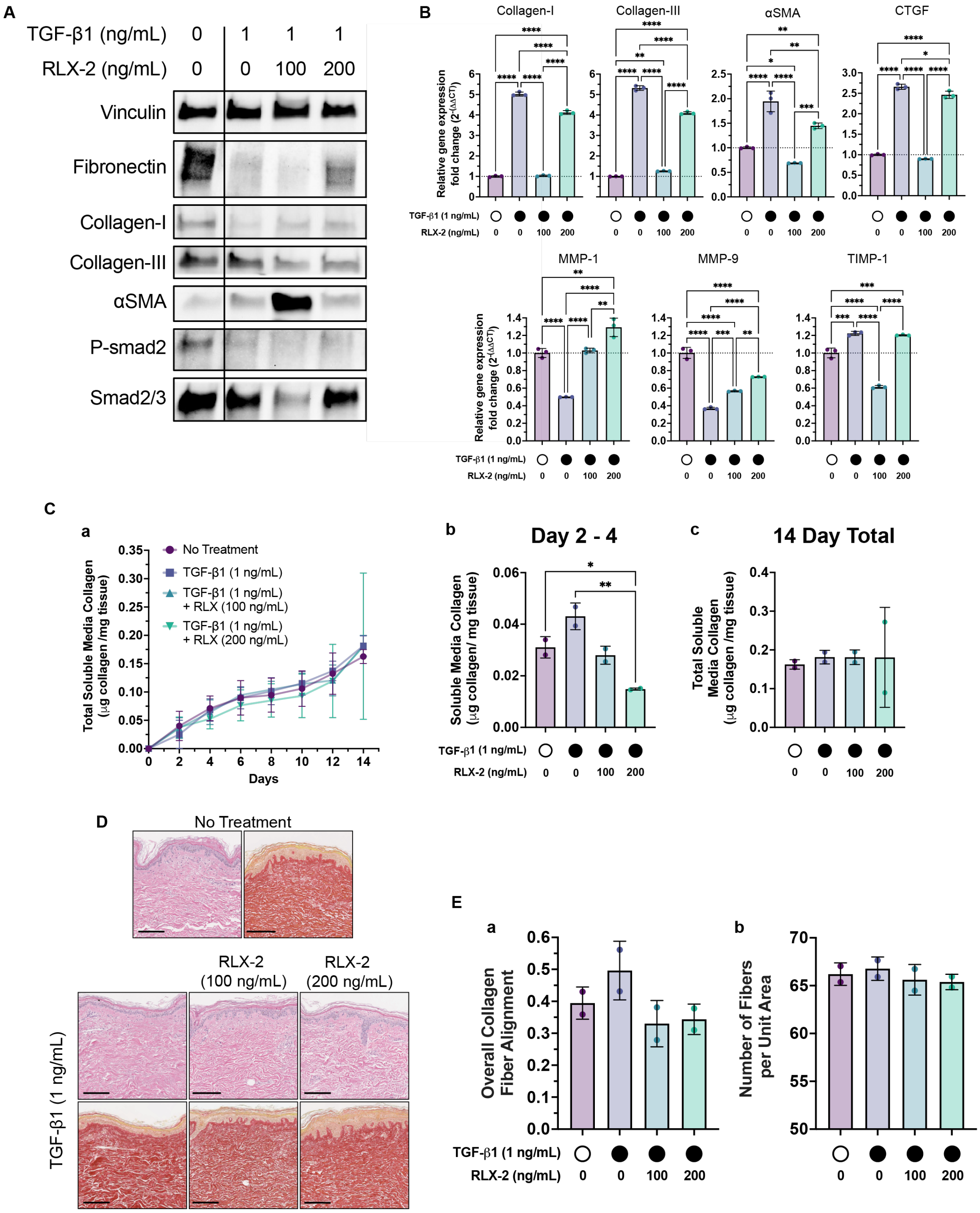


***Figure S1. RLX-2 inhibits the fibrotic scar phenotype in ex vivo normal human skin tissues.*** All data shown is using tissue from Patient 1 cultured *ex vivo* with TGF-β1 (1 ng/mL) with and without RLX-2 (100 or 200 ng/mL) except in (E). (**A**) Protein expression of ECM proteins and P-smad2. (**B**) RT-qPCR of ECM proteins (collagen-I, collagen- III, αSMA, CTGF) and ECM remodeling proteins (MMP-1, MMP-9, TIMP-1) implicated in wound healing and scar formation. (**C**) (a) Cumulative soluble collagen released in the media every two days over 14 days. (b) Soluble collagen released in media between treatment days 2 and 4. (c) Total soluble collagen released after 14 days. Collagen content was normalized to the total mass of the biopsy punch. Bars show range of biological duplicates (n=2 biopsy punches from the same patient). (**D**) Representative images of H&E and Sirius red staining of normal human skin tissues after 14 days of *ex vivo* culture with and without TGF-β1 and RLX-2 treatment. Top = H&E, Bottom = Sirius red. Scale bar = 200 µm. (**E**) CurveAlign analyses on (a) overall fiber alignment and (b) fiber density from Sirius red stained images of skin tissues. Points represent average of analysis performed on 8 ROIs per tissue section. Bars show range of averages from 2 different patient samples/ experiments (n=8 ROIs, N=2 patients). Statistical significance was determined by an ordinary one-way ANOVA using a Tukey test to control for multiple comparisons. *=p<0.05, **=p<0.005, ****=p<0.0001.

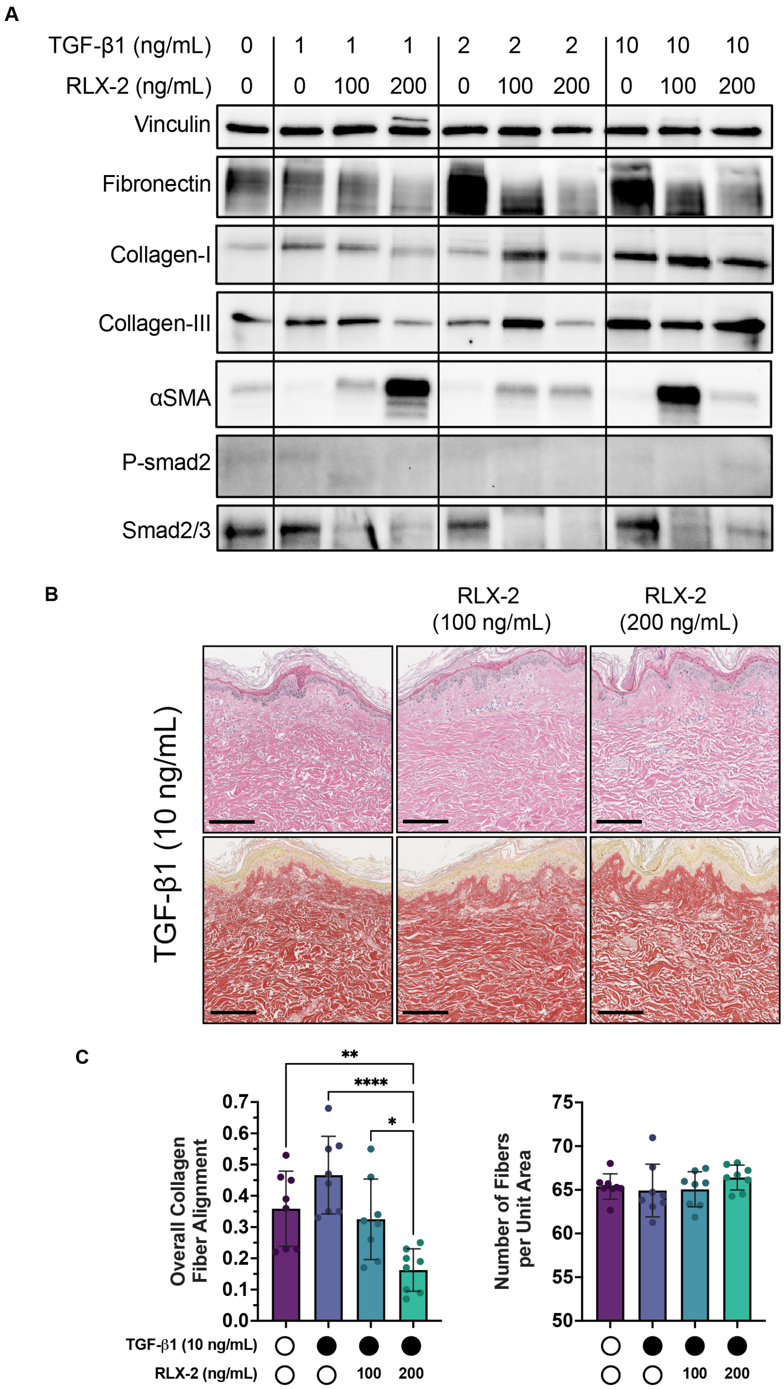


***Figure S2: Phenotypic changes in normal skin tissue cultured ex vivo from Patient 2.*** All data shown is using tissue from Patient 2 cultured *ex vivo* with TGF-β1 (1 – 10 ng/mL) with and without RLX-2 (100 or 200 ng/mL). (**A**) Protein expression of ECM proteins and P-smad2. (**B**) Representative ROIs of skin tissues treated with TGF-β1 (10 ng/mL) and RLX-2 (100 and 200 ng/mL) for 14 days. Top = H&E, Bottom = Sirius red. Scale bar = 200 μm. (**C**) CurveAlign analyses of overall fiber alignment (Left) and fiber density (Right) of normal human skin tissues cultured with TGF-β1 (10 ng/mL) with and without RLX-2 (100 or 200 ng/mL). Error bars show standard deviation of 8 ROIs from a single tissue section from Patient 2 (n=8 ROIs, N=1 patient). All analyses were performed on Sirius red stained sections. Statistical significance was determined by an ordinary one-way ANOVA using a Tukey test to control for multiple comparisons. *=p<0.05, **=p<0.005, ****=p<0.0001.

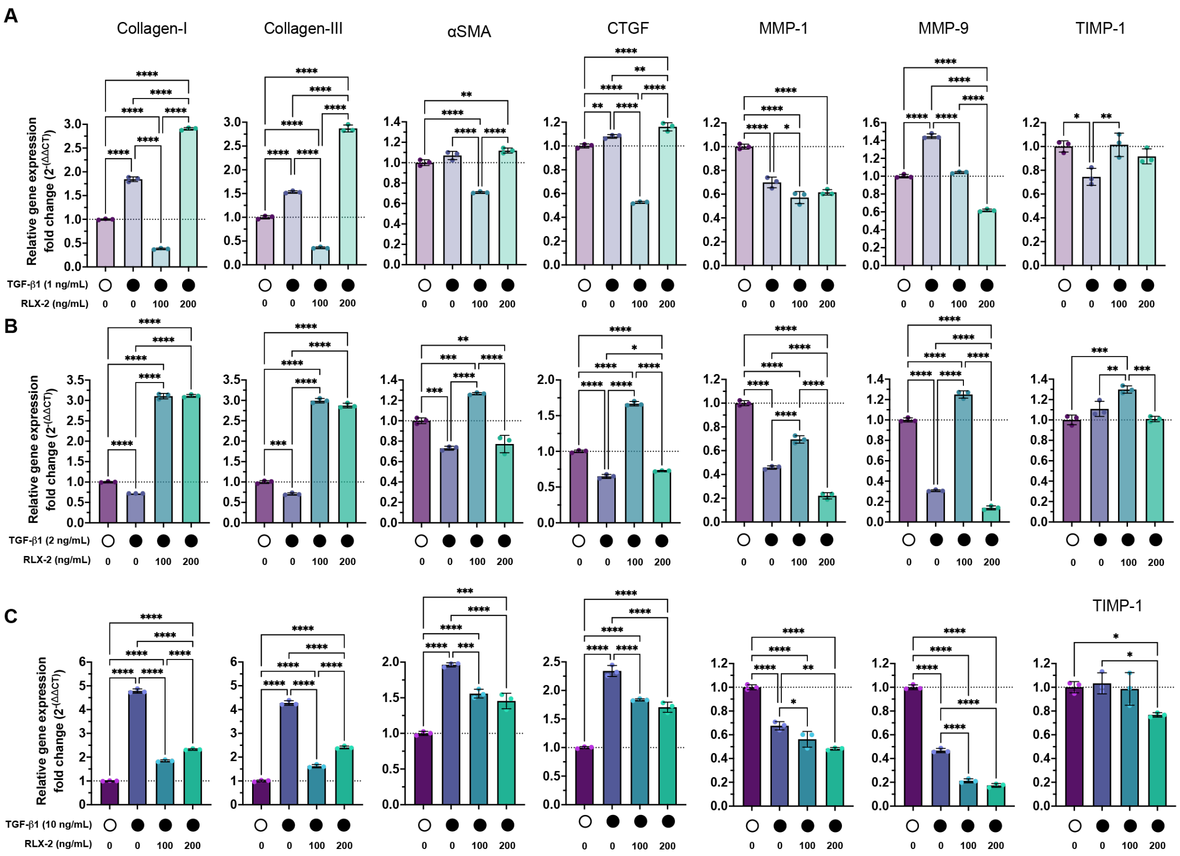


***Figure S3: Gene expression changes in ex vivo normal human skin tissues from Patient 2.*** All data shown is using tissue from Patient 2 cultured *ex vivo* with TGF-β1 (1 – 10 ng/mL) with and without RLX-2 (100 or 200 ng/mL) for 14 days. RT-qPCR of ECM proteins (collagen-I, collagen- III, αSMA, CTGF) and ECM remodeling proteins (MMP-1, MMP-9, TIMP-1) implicated in wound healing and scar formation. Normal skin tissues were treated with (**A**) TGF-β1 at 1 ng/mL, (**B**) TGF-β1 at 2 ng/mL and (**C**) TGF-β1 at 10 ng/mL with and without RLX-2 (100 or 200 ng/mL). Relative gene expression fold change is compared to B2M as the reference gene and the untreated control group in each set. Error bars show standard deviation of technical replicates from RT-qPCR. Statistical significance was determined by an ordinary one-way ANOVA using a Tukey test to control for multiple comparisons. *=p<0.05, **=p<0.005, ***=p<0.0005, ****=p<0.0001

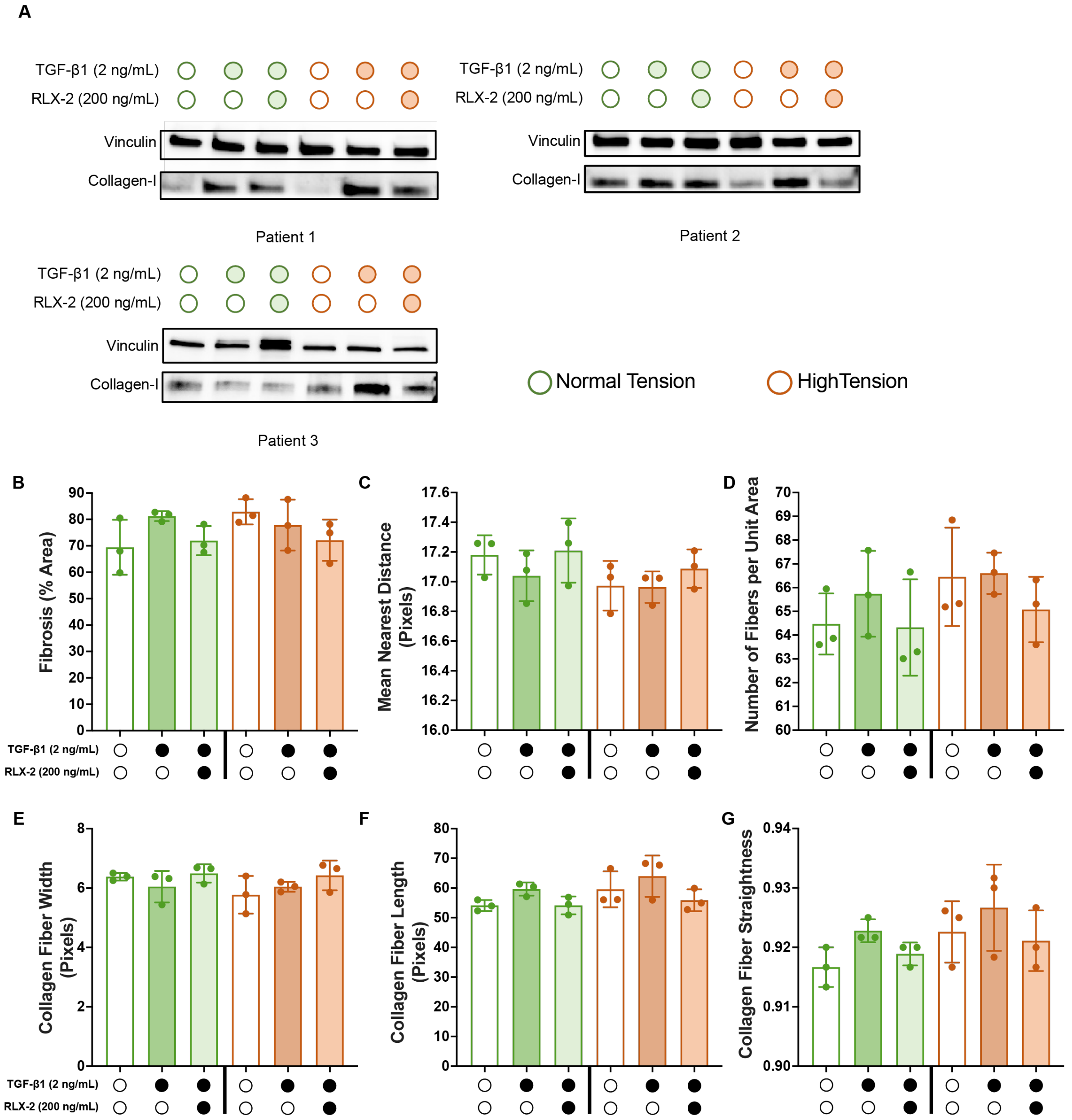


***Figure S4: Histological assessments of TenSkinTM tissues.*** (**A**) Collagen-I protein expression for each of three patient tissue samples. (**B**) Fibrosis quantification from Trichrome stained tissue sections using ImageJ. CurveAlign analyses of (**C**) mean nearest distance and (**D**) fiber density from Sirius red stained tissue sections. CT Fire analyses of collagen fiber (**E**) width, (**F**) Length and (**G**) straightness from Sirius red stained tissue sections. Each point represents the average of analyses performed on 6 ROIs per tissue section per patient. Three different patient tissue samples were assessed (n = 6 ROIs, N=3 patients). Bars outlined in green indicate skins under normal tension and bars outlined in orange indicate skins under high tension. Statistical significance was determined by an ordinary one-way ANOVA using a Tukey test to control for multiple comparisons. No significance was found between treatment groups, but the data trend towards changes in the collagen fiber architecture with treatments.

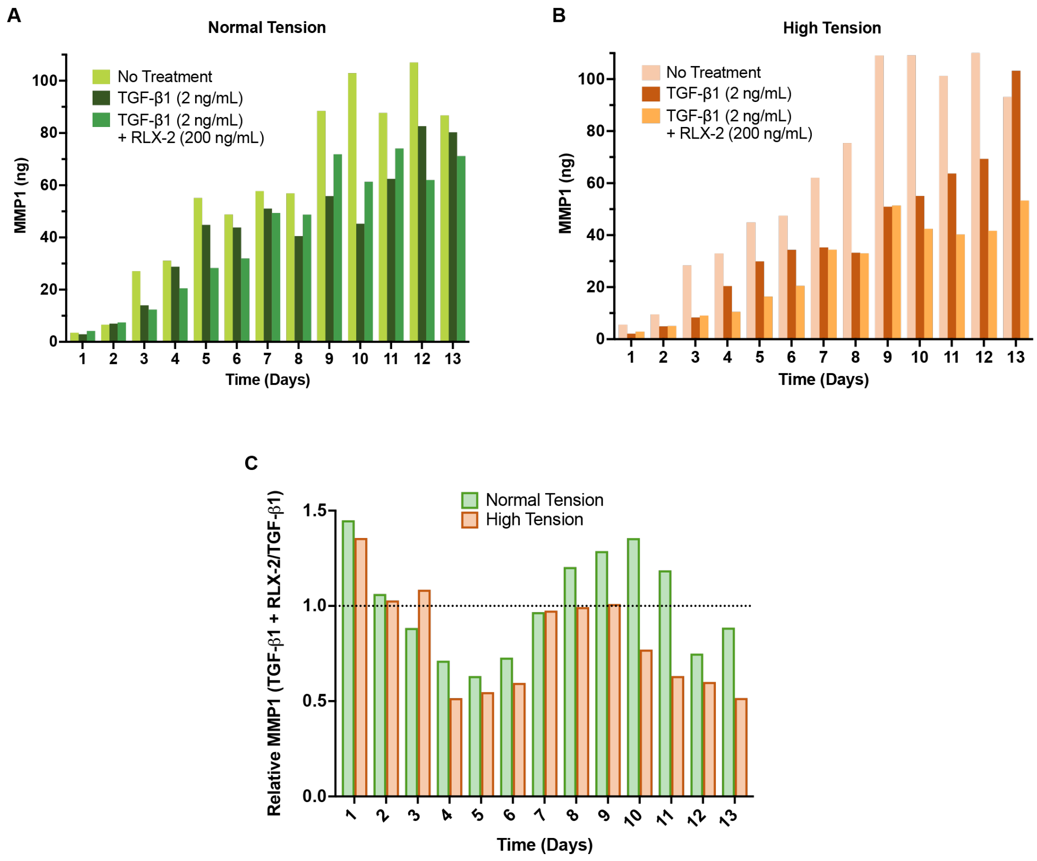


***Figure S5: MMP-1 activity in ex vivo TenSkinTM tissues.*** Endogenous active MMP-1 enzymes were measured in media samples collected from the TenSkinTM models every day during *ex vivo* culture. (**A, B**) MMP-1 each day in the media from (**A**) skin tissues under normal tension and (**B**) skin tissues under high tension and treated with TGF-β1 (2 ng/mL) with and without RLX-2 (200 ng/mL) for 14 days. (**C**) The ratio of MMP-1 in media from tissues treated with TGF-β1 (2 ng/mL) and RLX-2 (200 ng/mL) versus TGF-β1 (2 ng/mL) alone each day. Green bars indicate skins under normal tension and orange bars indicate skins under high tension.

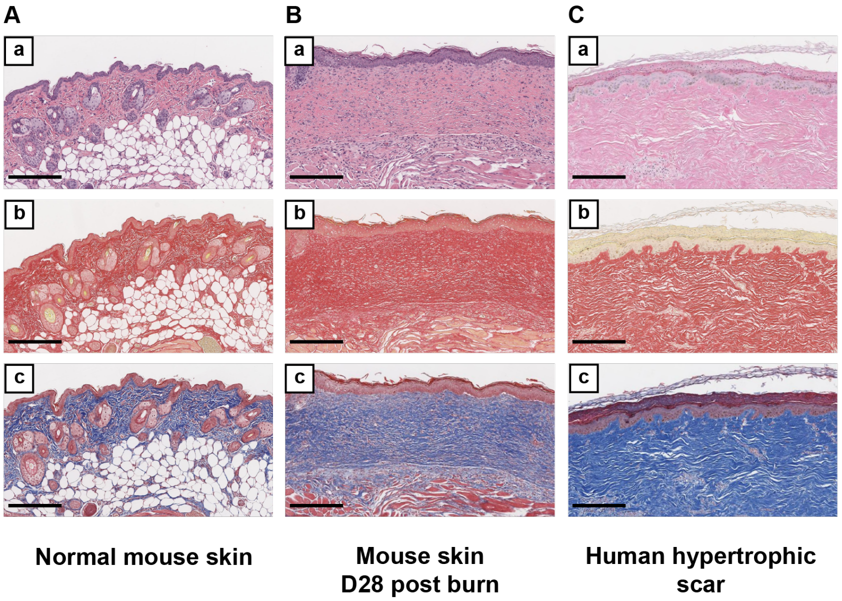


***Figure S6: Histological comparison of murine hypertrophic scar with human hypertrophic scar.*** (**A**) Normal mouse skin. (**B**) Scar created on mouse skin with 2N NaOH burn after healing 28 days. (**C**) Human hypertrophic scar. (a) H&E (b) Sirius Red, Collagen = red (c) Trichrome, Collagen = blue. Scale bar = 250 μm

**Supplementary Tables**

| Score | Collagen Fiber Appearance |
|---|---|
| 0 | normal |
| 1 | collagen is fine and fibrillar |
| 1.5 | Combination of 1 and 3 with highest percentage of fibers having the features of score 1 |
| 2 | Combination of 1 and 3 |
| 2.5 | Combination of 1 and 3 with highest percentage of fibers having the features of score 3 |
| 3 | collagen is fibrotic, vessels perpendicular to epidermis |
| 3.5 | combination of 3 and 5 with highest percentage of fibers having the features of score 3 |
| 4 | combination of 3 and 5 |
| 4.5 | combination of 3 and 5 with highest percentage of fibers having the features of score 5 |
| 5 | collagen is extremely sclerotic and compacted in thick bundles |

**Table S1: Pathologist scoring criteria for overall collagen fiber assessment in skin tissue sections.**

| Score | Collagen Fiber Orientation |
|---|---|
| 0 | normal basket-weave pattern |
| 1 | <25% abnormal |
| 2 | 26-50% abnormal |
| 3 | 51-75% abnormal |
| 4 | 76-100% abnormal |
| 5 | keloid-like fiber orientation |

**Table S2: Pathologist scoring criteria for collagen fiber orientation assessment in skin tissue sections.**

| Score | Collagen Fiber Density |
|---|---|
| 0 | normal fiber bundle density |
| 1 | <25% abnormal |
| 2 | 26-50% abnormal |
| 3 | 51-75% abnormal |
| 4 | 76-100% abnormal |
| 5 | keloid-like fibers |

**Table S3: Pathologist scoring criteria for collagen fiber density assessment in skin tissue sections.**

| Score | Collagen Fiber Maturity |
|---|---|
| 0 | normal fiber maturity |
| 1 | <25% abnormal fibers |
| 2 | 26-50% abnormal fibers |
| 3 | 51-75% abnormal fibers |
| 4 | 76-100% abnormal fibers |
| 5 | keloid-like fibers |

**Table S4: Pathologist scoring criteria for collagen fiber maturity assessment in skin tissue sections.**
